# Tropism of SARS-CoV-2 in commonly used laboratory cell lines and their proteomic landscape during infection

**DOI:** 10.1101/2020.08.28.271684

**Authors:** Elisa Saccon, Xi Chen, Flora Mikaeloff, Jimmy Esneider Rodriguez, Laszlo Szekely, Beatriz Sá Vinhas, Shuba Krishnan, Siddappa N. Byrareddy, Teresa Frisan, Ákos Végvári, Ali Mirazimi, Ujjwal Neogi, Soham Gupta

## Abstract

The present pandemic caused by severe acute respiratory syndrome coronavirus-2 (SARS-CoV-2) is driving intense research activities to understand the basic biology of the virus and determine effective therapeutic strategies. The commonly used laboratory cell lines of human origin are the first line of experimental models to study the pathogenicity and performing antiviral assays. Thus, to find suitable cell models to study SARS-CoV-2, we assessed the tropism and cytopathogenicity of the first Swedish isolate of SARS-CoV-2 in six different cell lines of human origin and compared their growth characteristics to other globally isolated strains. Overall, Calu-3, Caco2, Huh7, and 293FT cell lines showed a high to moderate level of susceptibility to the majority of virus isolates. In Caco2 cells the virus can achieve high titers in the absence of any prominent cytopathic effect. The protein expression profile during SARS-CoV-2 infection revealed cell-type-specific regulation of cellular pathways. Type-I interferon signaling was identified as the common dysregulated cellular response in Caco2, Calu-3 and Huh7 cells. Overall, cell-type specific variability was noted for cytopathogenicity, susceptibility and cellular response to SARS-CoV-2. This study provides important clues regarding SARS-CoV-2 pathogenesis and can represent as a guide for future studies to design therapeutics.

## Introduction

Severe acute respiratory syndrome coronavirus 2 (SARS-CoV-2), the causative agent of coronavirus disease 2019 (COVID-19) pandemic, is a highly pathogenic coronavirus that has created a global public health challenge (Hu et al., 2020). The virus primarily attacks the lung and the patients often present with severe respiratory distress. However, apart from respiratory symptoms, involvement of other organs with cardiovascular, gastrointestinal, liver, neurological, hematological and skin manifestations in the disease pathology has been documented, suggesting the vulnerability of these anatomical sites to this virus (Gavriatopoulou et al., 2020; Trypsteen et al., 2020). This has been attributed to the presence of the primary receptor of the virus, angiotensin-converting enzyme 2 (ACE2), throughout the body (Hamming et al., 2004). However, we still lack information on the basic virology and the pathogenesis of the virus in different organs.

In the current pandemic, numerous research activities have been focusing on understanding the mechanisms of viral pathogenesis and developing effective antivirals against the infection (Appelberg et al., 2020; Bojkova et al., 2020; Chu et al., 2020; Jureka et al., 2020; Zecha et al., 2020). In this scenario, Vero-E6 cell-line has been widely employed for viral isolation, propagation and antiviral testing, due to its high virus production and a prominent cytopathic effect (CPE) upon infection. However, since Vero-E6 originates from monkey and lacks the gene cluster associated with type-I interferon (IFN-I), a more physiologically relevant cell-line of human origin is warranted, that is able to better recapitulate the complexity of host cellular response to the SARS-CoV-2 in humans. Furthermore, the use of cell lines originating from different organs of the body to certain extent can reflect the organ-specific response to the infection and may provide a better understanding of the pathogenesis and transmissibility within body and aid in designing host-targeted therapeutics.

In this study, we aimed to determine the SARS-CoV-2 tropism in six commonly used laboratory human cell lines that originate from the lungs, intestine, liver and kidney. Our data provide a comprehensive view of how SARS-CoV-2 behaves differently in different human cell lines depending on the viral strain, the cell line susceptibility and, possibly, the isolation method. Overall, the data presented in this manuscript will be beneficial for choosing a suitable cell system to investigate viral pathogenicity and development therapeutics against SARS-CoV-2.

## Results

### Susceptibility and cytotoxicity of first Swedish SARS-CoV-2 isolate in commonly used laboratory cell lines

We infected Vero-E6, Calu-3, A549, Caco2, Huh7, 293FT and 16HBE with the first Swedish isolate of SARS-CoV-2 virus (SWE/01/2020) at a multiplicity of infection (moi) of 1 and 0.1 as previously described (Appelberg et al., 2020). Virus-induced cytotoxicity was evaluated by measuring the cellular ATP using Viral-ToxGlo™ assay (Promega) and virus production was determined by measuring the presence of viral genome in the cell culture supernatant using qPCR to determine the N-gene RNA (Corman et al., 2020) starting at 3h post-infection (hpi) and followed up to 120hpi (Appelberg et al., 2020). As shown in Figure 1A, infection with moi 0.1 or 1 showed a very similar pattern of virus production over time and by the end of 120h attained similar viral copies in the supernatant. Infection at moi 0.1 induced significant cytopathogenicity in Vero-E6 (less than 3% viability by 48hpi) and a significant increase of 3.8 log10 viral RNA copies in the supernatant at 24hpi. Of the six human cell lines that were tested, Caco2 (intestinal) and polarized Calu-3 (lung) showed the highest virus production with >4 log10 RNA copies by 48hpi (*p<0.001*) and thereafter marginal increase till 120hpi. It was interesting to note that Calu-3 cells, which were infected after 72h of seeding and showed tightly closed together cells with polygonal or cuboidal features and defined boundary, had a higher susceptibility to SARS-CoV-2 compared to Calu-3 cells that were infected after 24h of seeding (round and isolated). The enhanced susceptibility of Calu-3 with longer incubation of 72h prior to infection was possibly due to polarization of the cells (Foster et al., 2000), as was reported previously for SARS-CoV (Tseng et al., 2005). 293FT (kidney; *p<0.01*) and Huh7 (liver; *p<0.02*) showed moderate virus production with >1 log10 viral RNA copies in the supernatant by 120hpi. 16HBE (lung) and A549 (lung) cells showed very poor virus production with <0.6 log10 RNA copies. Interestingly, other than Vero-E6, viral-induced cytotoxicity was only observed in Calu-3 cells with a viability of ≤50% by 48hpi and ≤80% by 72hpi. None of the other cell lines showed any apparent cytotoxicity (viability>85%) (Figure 1B). We observed that the virus production in susceptible cell lines reached saturation by 48hpi. Therefore, we investigated the changes caused in the cell surface of Calu-3, Caco2, Huh7, and 293FT cells during virus production at 48hpi (moi 0.1) using scanning electron microscopy (SEM). We did not observe any significant changes in the morphology of the cell surface in the mock-infected cells. In SARS-CoV-2 infected Calu-3 and Caco2 cells, numerous virus-like particles corresponding to the size of SARS-CoV-2 (approx. 70nM) were observed to be attached to the cell surface or cellular projections (Figure 2A). Interestingly, even though there was moderate virus production in Huh7 and 293FT cells we did not observe any attached virus-like particles on the cell surface after scanning several fields (data not shown). The possible reasons for this could be either low level virus production in Huh7 cells and 293FT cells compared to Calu-3 and Caco2 cells that was missed visually or the virus release mechanism is different than the budding out of the virus (Ghosh et al., 2020).

**Figure 1.**
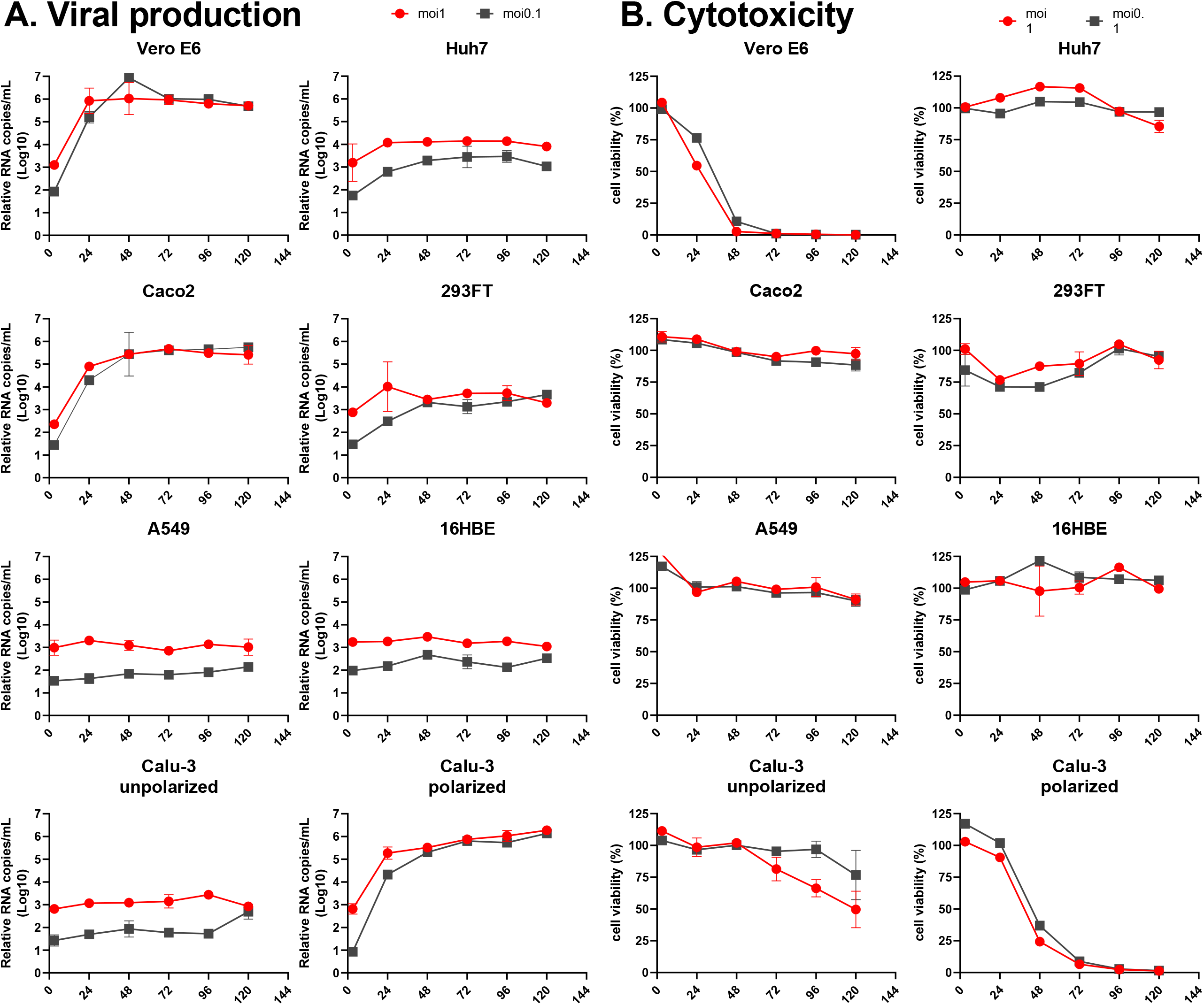
Viral production and cytopathogenicity of SARS-CoV-2 in Vero E6 and six different cell lines of human origin. Indicated cell lines were infected with SARS-CoV-2 at moi of 1 and 0.1 either in duplicate or triplicate. A. Viral supernatant samples were harvested at 3hpi, 24hpi, 48hpi, 72hpi, 96hpi and 120hpi. Viral production was determined by quantitative qRT-PCR targeting the N gene of SARS-CoV-2 comparing each time points to 3hpi. B. Cell viability was measured at 3h post infection (hpi), 24hpi, 48hpi, 72hpi, 96hpi and 120hpi by Viral-ToxGlo™ assay. The viability at each time was determined in comparison to the uninfected control.

**Figure 2.**
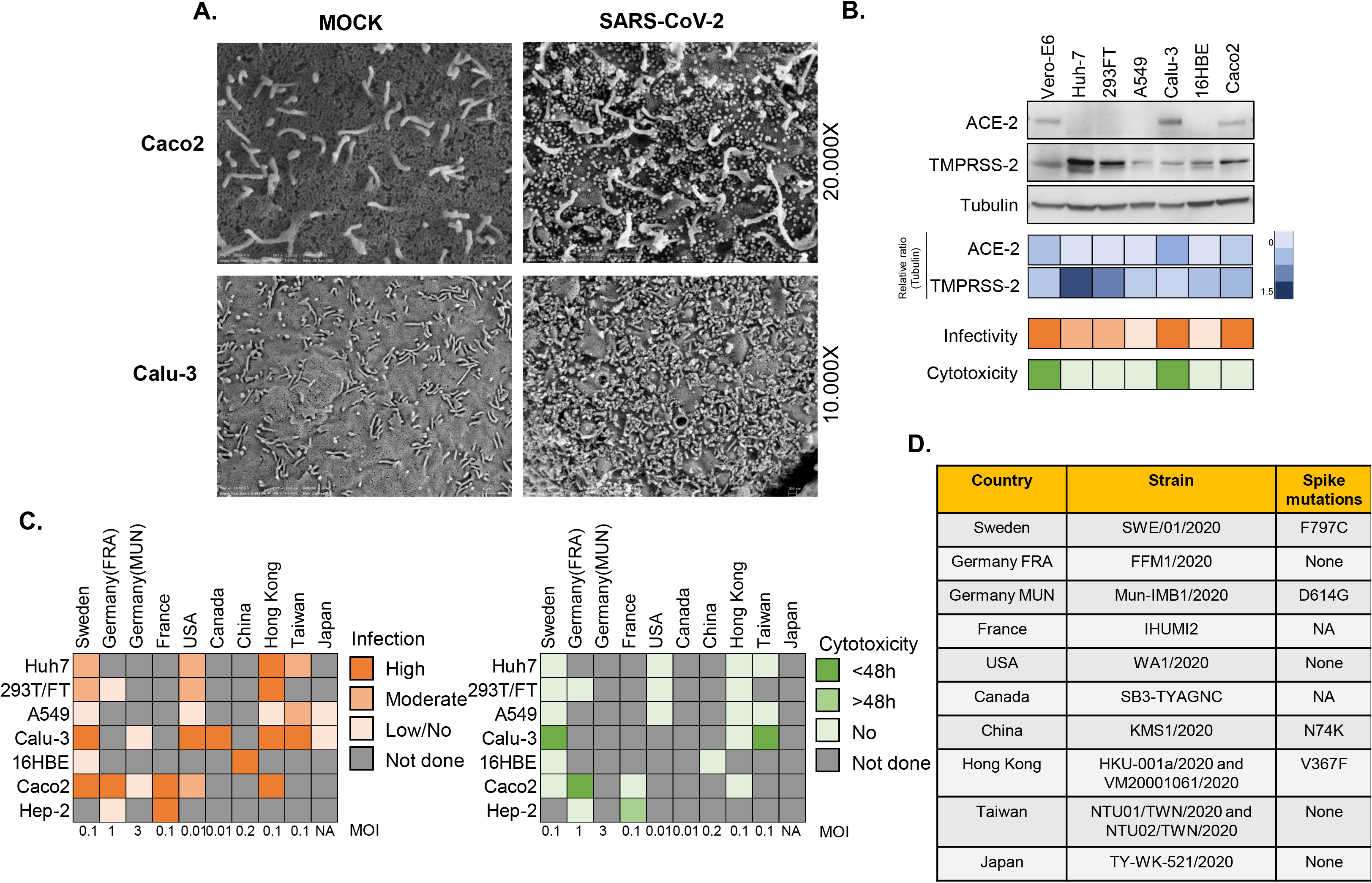
Cell-specific and strain-specific differences in SARS-CoV-2 tropism. A. Scanning electron microscopy images of the cell surface of mock-infected and SARS-CoV-2 infected (moi 0.1) Caco2 and Calu-3 cells. Indicated cell lines were infected with SARS-CoV-2 at an moi of 0.1 and fixed 48hpi for microscopy analysis. B. ACE2 and TMPRSS2 receptor expression in six human cell lines. ACE2 and TMPRSS2 expression in these cell lines were determined by western blot in the cell lysates. Heatmaps with relative protein quantification (blue), susceptibility (orange) and cytotoxicity (green) are shown. C. Susceptibility of globally isolated SARS-CoV-2 strains. Based on previously published or preprint articles, a chart was created showing the susceptibility of the commonly used laboratory cell lines and cytopathogenicity of SARS-CoV-2 strains isolated from different geographical locations. Boxes are color coded based on intensity as indicated in the figure. The moi used wherever available is mentioned in the bottom. D. Table showing presence of any amino acid substitution in the spike protein of the indicated strains. NA= Not available.

### Expression of ACE2 and TMPRSS2 in different cell lines

ACE2 receptor and transmembrane serine protease 2 (TMPRSS2) activity have been shown to be critical for SARS-CoV-2 entry into the cell (Jureka et al., 2020; Matsuyama et al., 2020). To correlate the ACE2 and TMPRSS2 expression with the tropism, we determined the protein expression in the cell lysates by western blot (Figure 2B). Among the infected cell lines, ACE2 expression was only observed in Vero-E6, Calu-3 and Caco2 that correlated with high virus production. When we looked into co-expression of ACE2 and TMPRSS2, it was most prominent in Caco2, followed by Vero-E6 and Calu-3, although Calu-3 showed low TMPRSS2 expression. Contrarily, Huh7 and 293FT strongly expressed TMPRSS2 but lacked ACE2 expression indicating that each receptor has an individual role in aiding the infection.

### Susceptibility and cytopathogenicity of globally isolated SARS-CoV-2 strains

To compare the tropism of the Swedish SARS-CoV-2 virus with the other globally isolated strains, we performed a literature survey to determine the susceptibility and cytopathogenicity in all the six cell lines as noted above. We have included virus isolates from Germany (FFM1/2020 (Bojkova et al., 2020) and Mun_IMB1/2020 (Zecha et al., 2020)), France (IHUMI2) (Wurtz et al., 2020), USA (WA1/2020) (Jureka et al., 2020), Canada (SB3-TYAGNC) (Banerjee et al., 2020), China (KMS1/2020) (Liao et al., 2020), Hong Kong (HKU-001a/2020 (Chu et al., 2020) and VM20001061/2020 (Hui et al., 2020), Taiwan (NTU01/2020 and NTU02/2020) (Hsin et al., 2020) and Japan (TY-WK-521/2020) (Matsuyama et al., 2020) and found major differences in cellular tropism and cytopathogenicity (Figure 2C). Specifically, we observed that the majority of the strains were able to infect both Caco2 and Calu-3, except for the Muc_IMB1/2020 (both the cell lines not susceptible) (Zecha et al., 2020) and the Japanese strain (Calu-3 cells not susceptible) (Matsuyama et al., 2020). The Frankfurt_FFM1/2020 strain showed a rapid CPE in Caco2 by 24hpi at moi 0.1 (Bojkova et al., 2020), while other strains, including the Swedish isolate, did not show any prominent CPE either at higher infective dose or with prolonged time of incubation (Chu et al., 2020). Additionally, using the Wuhan/Hu-1/2019 strain as reference, we compared the amino acid changes in the spike protein of these strains. The Frankfurt and Taiwan strains showed no dissimilarities, while the Swedish (F797C), Munich (D614G), China (N74K) and Hong Kong (V367F) strains each showed a single amino acid substitution (Figure 2D).

### Proteomic analysis of the cell lines

Since we observed differential susceptibility to SARS-CoV-2 infection in different cell lines originating from the human lung (Calu-3), intestine (Caco2), liver (Huh7) and kidney (293FT), we investigated how the cellular proteins are regulated during infection in these cell lines. To this end, we either infected or mock-infected polarized Calu-3 cells, Caco2 cells, Huh7 cells and 293FT cells with SARS-CoV-2 (moi 1) in triplicates. The cells were harvested 24hpi, lysed and equal concentration of the protein was used to perform quantitative proteomics using a TMT-labeling strategy as previously described (Appelberg et al., 2020). Among the four cell lines, Calu-3 showed major changes in protein abundance upon infection, with 6462 proteins differentially expressed in infected cells than the mock, followed by Caco2 with a significant difference in 177 proteins. No change in the global protein abundance was observed in Huh7 (only four proteins differentially expressed) and 293FT (no proteins differentially expressed) at 24hpi (Figure 3A, Supplemental Table S3). The virus secretion in the cell culture supernatant at 3hpi and 24hpi is shown in Figure 3B. The viral protein abundance in the cells is shown in Figure 3C. The proteins that were detected are ORF3a, ORF6, ORF7a, ORF8, ORF9b, M, N, S and Replicase polyprotein 1ab (abbreviated as rep). The higher abundance of viral proteins detected in Calu-3 correlated with higher level of virus production and the change in host protein abundance.

**Figure 3.**
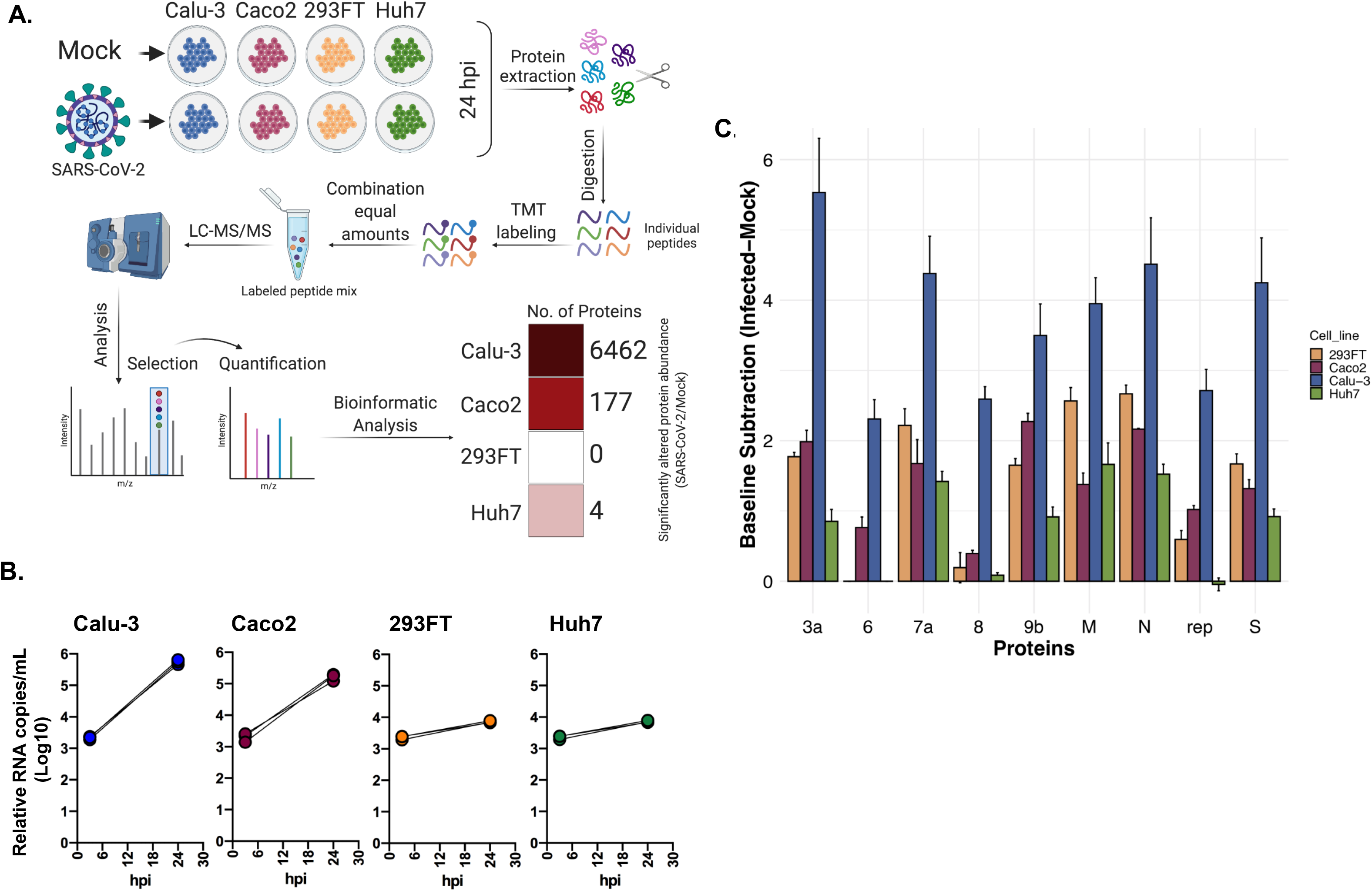
Proteomic landscape of different commonly used laboratory cell lines upon SARS-CoV-2 infection. A. Brief methodology of the proteomics experiment in the indicated cell lines that were either mock-infected or infected with SARS-CoV-2 moi 1 for 24h. Significantly altered host proteins in the indicated cell lines are represented in the heatmap. B. Relative viral production in the cell supernatant, measured at 3hpi and 24hpi. C. Detected viral proteins in the indicated cells by tandem mass tag-labeled mass spectrometry (TMT-MS).

### Type-I IFN signaling is commonly dysregulated in Calu-3, Caco2 and Huh7 cell lines by SARS-CoV-2

Since Calu-3 and Caco2 were the only cell lines that showed substantial protein regulation upon infection at 24h, we compared the significantly regulated proteins in those two cell lines. As shown in the Venn diagram in Figure 4A, there were 132 proteins that were commonly dysregulated in both Calu-3 and Caco2. Among the 132 dysregulated proteins, 88 proteins were similarly upregulated (44 proteins) and downregulated (44 proteins) in both the cell lines (Supplemental figure S1). Reactome pathway analysis on the 132 significantly altered proteins common to both cell lines showed a strong enrichment of type-I and type-II interferon signaling pathways and its related RIG-I/MDA5 (DDX58/IFIH1) signaling pathway (Figure 4B). To investigate which are the proteins associated with this pathway changing upon infection, as a next step, we assessed the changes induced by the infection in the levels of each protein associated with the following pathways: interferon response, including the interferon alpha/beta signaling (Pathway: R-HSA-909733), interferon gamma signaling (Pathway: R-HSA-877300) and antiviral mechanism by IFN-stimulated genes (ISGs, Pathway: R-HSA-1169410). As shown in the heatmaps, 105 (out of 129 detected) and 27 (out of 131 detected) were differentially regulated in Calu-3 (Figure 4C, Supplemental Table S4), and Caco2 respectively (Figure 4D, Supplemental Table S4). At 24hpi, both 293FT and Huh7 did not show any differentially regulated protein belonging to IFN-signaling pathways (109 detected proteins; Supplemental Table S4) and the heat maps are shown in Supplemental Figure S2. As shown in the heatmaps, among 129 detected proteins belonging to these pathways, 105 were differentially regulated in Calu-3 (63 upregulated and 42 downregulated), while in Caco2 among 131 detected proteins 27 were differentially regulated (25 upregulated and 2 downregulated) (Figure 4C and 4D, Supplemental Table S4). The protein-protein interaction network of the significantly altered proteins showed two definite clusters in Calu-3: one including proteins associated with RIG-I (DDX58) and Type-I/II signaling complex and another majorly including components of nucleoporin complex that were down-regulated. The karyopherin family, and a single cluster (RIG-I (DDX58) and Type-I signaling complex) were upregulated in Caco2 (Supplemental Figure S3A and S3B). In general, we observed an interferon stimulation in SARS-CoV-2 infected Calu-3 and Caco2 cells and SARS-CoV-2 receptor ACE2 has been considered to be an interferon stimulatory gene (Ziegler et al., 2020). However, we did not observe any significant differences in the protein levels of ACE2 or TMPRSS2 upon infection in our proteomics data, rather ACE2 was down-regulated in SARS-CoV-2 infected cells (Supplemental Figure S4).

**Figure 4.**
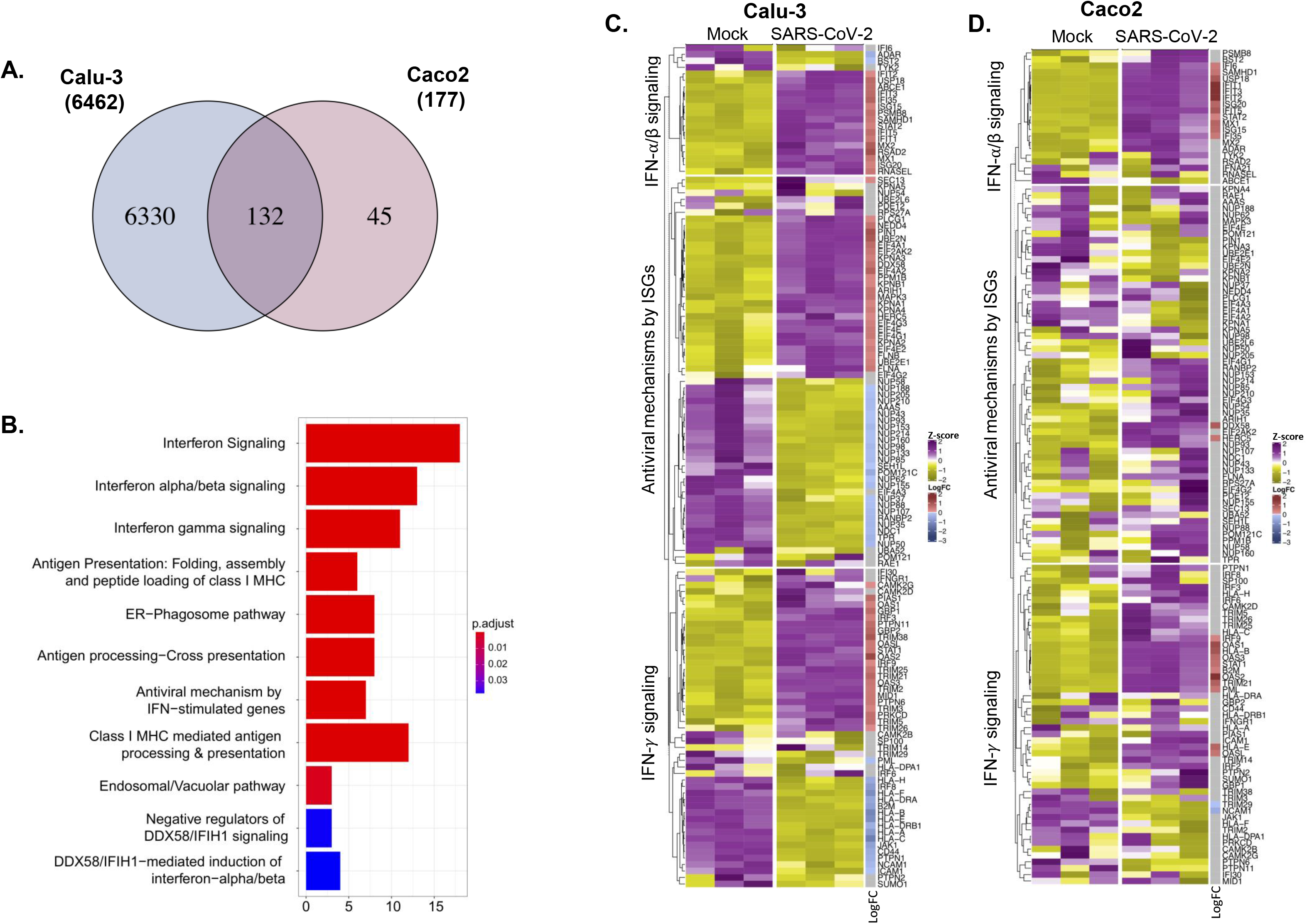
Interferon-signaling pathways are commonly dysregulated by SARS-CoV-2 infection in Caco2 and Calu-3 cell lines. A. Venn diagram and overlap of proteins with significant abundance between mock and 24hpi in Caco2 and Calu-3 cell lines. B. Barplot enrichment map of significant overlapping proteins in Caco2 and Calu-3 cell lines from ReactomePA. Number of proteins are indicated on the x axis. P values are indicated from highly significant in red to significant in blue. C. Heatmap representing the number of significant proteins (LIMMA, FDR < 0.05) between mock and 24hpi in Calu-3 cell line. D. Heatmap representing the number of significant proteins (LIMMA, FDR < 0.05) between mock and 24hpi in Caco2 cell line.

Type-I IFN, like IFN-β, plays a major role in host defenses against various viruses and in our proteomics data we observed dysregulation of type-I IFN response associated proteins including ISGs, therefore we assessed the mRNA expression of IFN-β and the downstream ISGs (IFIT1, MX1, MX2, ISG15 and RIG-I) using qPCR. In line with our proteomics findings, the mRNA levels of IFN-β and all the tested ISGs significantly increased upon SARS-CoV-2 infection in Calu-3 and Caco2 cells. 293FT and Huh7 cells did not show any significant variations at 24hpi (Figure 5A). IFN-β is released by cells through a signaling cascade that is initiated after activation of RIG-I or RLRs upon interaction with viral products. The western blot analysis showed a major increase in activation of RIG-I, MDA-5 and its downstream effectors p-IRF3, p-STAT1 and ISG15 in infected Calu-3 and Caco2 cells (Figure 5B and 5C) corroborating the qPCR results. In general, the steady-state level of RIG-I, MDA-5 and ISG15 were higher in Caco2 and Calu-3 cells as compared to 293FT and Huh7 cells and the latter two showing no comparable changes following 24hpi (Figure 5C). No observable change was noted in conjugated ISG15 in any of the cell lines (Supplemental Figure S5A and S5B). It is conceivable that the observed changes in SARS-CoV-2 infected Calu-3 and Caco2 cells 24hpi could be due to high level of infectivity as observed by the expression of SARS-CoV-2 nucleoprotein (Figure 5B and 5C) and higher susceptibility (Figure 1B and 3B) as compared to 293FT and Huh7.

**Figure 5.**
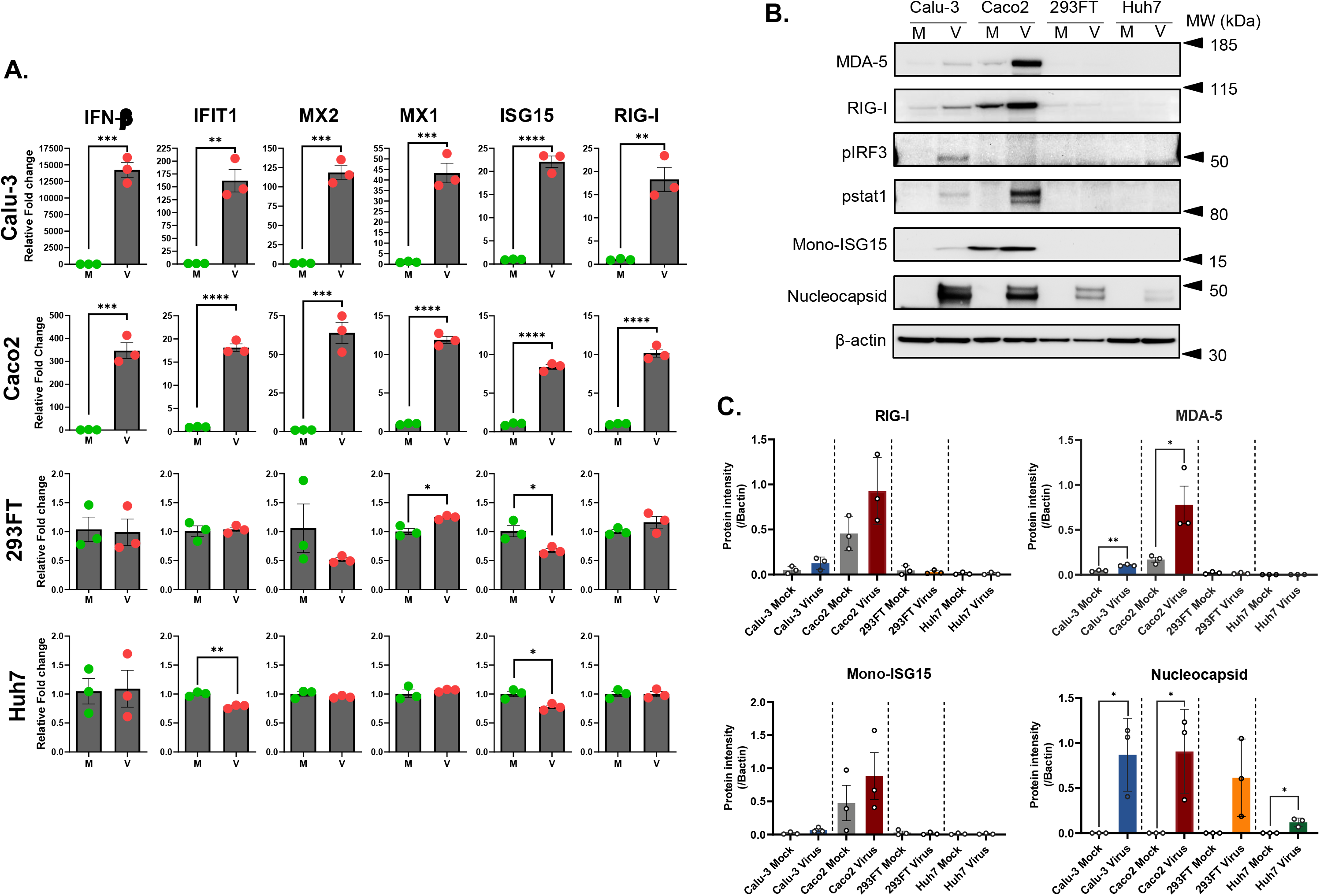
Different expression of ISGs in the infected cell lines. A. The gene expression level of IFN-β and indicated representative interferon stimulated genes (ISGs): IFIT1, MX2, MX1, ISG15 and RIG-I in Calu-3, Caco2, 293FT and Huh7 cell lines 24hpi. The results are shown as fold change relative to mock-infected cells, normalized to GAPDH. The mean ± SEM of at least two independent experiments is shown. M = mock; V = virus. B. Protein expression levels of MDA-5, RIG-I, pIRF-3, p-STAT1, and viral nucleocapsid in infected and mock-infected Calu-3, Caco2, 293FT and Huh7 cells 24hpi. C. The intensity of specific bands for MDA-5, RIG-I, mono-ISG15 and SARS-CoV-2 nucleoprotein was quantified by ImageJ and fold change was calculated relative to the mock-infected cells, normalized to β-actin. The intensity of p-IRF3 and p-STAT1 was not quantified as expression could be observed only in one of the experimental replicates. The mean ± SEM of at least two experiments is shown.

Previous proteomics-based studies have observed that SARS-CoV-2 causes global proteomic changes after 48hpi specifically in pathways related to ErbB, HIF-1, mTOR and TNF signaling (Appelberg et al., 2020), complement system and coagulation cascades (Tiwari et al., 2020) and interferon signaling (Unpublished data). In order to understand the delayed changes in Huh7 cells (Huh7^48h^) compared to Calu-3 and Caco2 cells, we extracted the proteomics data set from our earlier study (Appelberg et al., 2020). Next, we aligned the significantly changed proteins in this data set with that of the Calu-3 and Caco2 cells. There were 42 common proteins that were dysregulated in all three cell lines (Figure 6A), showing similar trend in 15 proteins (8 were commonly upregulated and 7 were commonly downregulated) and opposite trends in 27 proteins in all the three cell lines (Supplemental Figure S6). We employed Reactome pathway analysis to define the commonly dysregulated pathways using 42 proteins and observed that pathways related to interferon signaling were the top hits (Figure 6B). We looked in detail into the 24 proteins that belonged to pathways related to IFN-signaling and were significantly altered in Calu-3 and Caco2. As shown in Figure 6C, only STAT2, STAT1, DDX58, ISG15 and IFIT1 were commonly upregulated in all three cell lines. Distinct features were noticed in B2M, PML, HLA-E and HLA-B in Calu-3 (downregulated), Caco2 (upregulated) and Huh7^48h^ (no change). IFI35 was upregulated in both Calu-3 and Caco2 cells but was downregulated in Huh7^48h^.

**Figure 6.**
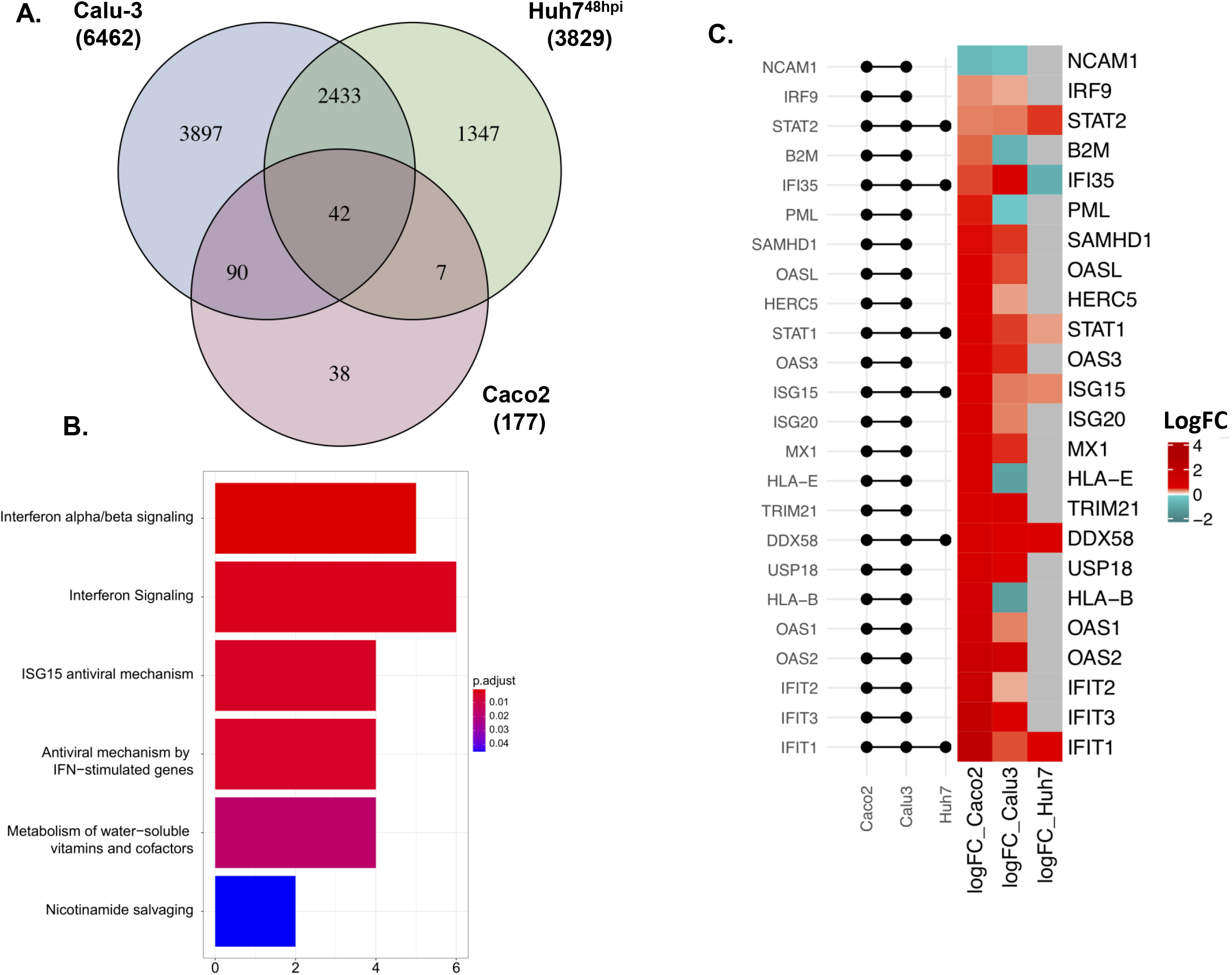
Overlap of three cell lines confirm the implication of interferon pathway in SARS-CoV-2 infection. A. Venn diagram and overlap of proteins with significant abundance between mock and 24hpi in Caco2, Calu-3 and Huh7 cell lines. B. ClusterProfiler enrichment map of significant proteins in Caco2, Calu-3 and Huh7 cell lines. Size of the bubble is proportional to gene ratio. C. ReactomePA barplot enrichment map of significant overlapping proteins in Caco2, Calu-3 and Huh7 cell lines.

### Cell-type and virus specific differences in proteome

There are obvious commonalities and distinct features between different cell lines that could be defined by the tissue of origin. Till now we have shown that SARS-CoV-2 commonly regulates the type-I IFN signaling in cell lines derived from different tissues. In order to determine which other pathways are distinctively regulated by SARS-CoV-2 in Calu-3, Caco2 and Huh7^48h^ cells, we identified the proteins that were significantly uniquely altered by infection in each of the 3 cell lines. There were 3897 proteins in Calu-3, 1347 proteins in Huh7^48h^ and 38 proteins in Caco2 that uniquely changed (Figure 6A). A Reactome pathway analysis on these proteins revealed major changes in pathways related to mitochondrial processes in Calu-3 (Supplemental Figure S7A), and eukaryotic translation processes in Huh7^48h^ (Supplemental Figure S7B). We could not identify clearly any specific pathway modulated in Caco2. Additionally, we investigated altered pathways shared between only Huh7^48h^ and Caco2 and only Huh7^48h^ and Calu-3. As presented in Supplemental Figures S8A and S8B respectively, unlike Caco2 and Calu-3 they did not show shared characteristics of IFN-signaling dysregulation as top pathways.

## Discussion

Cellular models that reproduce the SARS-CoV-2 life cycle are essential to understand the viral-host interplay and to test new antivirals. Vero-E6 cell-line that originates from African green monkey kidney cells is often a choice for cell culture-based infection model for coronavirus research (Ogando et al., 2020), since it expresses the receptor ACE2 and shows prominent cytopathogenicity with efficient virus production. However, Vero-E6 may not be a suitable cell model to study the pathophysiology of the cell in response to the virus infection as it lacks genes encoding type-I interferons (Osada et al., 2014). Cell lines of human origin are more relevant. In the present study, we systematically analyzed SARS-CoV-2 susceptibility and cytopathogenicity in cell lines originating from human lung (Calu-3, A549, 16HBE), colon (Caco2), liver (Huh7) and kidney (293FT). Furthermore, using proteomics we characterized the cellular changes caused by SARS-CoV-2 in the susceptible cell lines Calu-3, Caco2, 293FT and Huh7. Our data provides insight into the cell-type-specific cellular re-organization caused by SARS-CoV-2.

Several recent studies testing different local strains show conflicting results concerning susceptibility and cytopathogenicity in human cell lines (Figure 2C), mostly when Caco2 and Calu-3 cells are employed. The Swedish isolate tested here showed high virus production and prominent cytopathogenicity in Calu-3 cells. However, it needs to be pointed out that the susceptibility of the Calu-3 cells increased following polarization of the cells after 72h of incubation before infection, as with 24h of incubation of the cells prior to infection showed a very moderate virus production and a delayed CPE (Figure 1). Another observation with Calu-3 cells was that upon infection, they did not show a CPE similar to Vero-E6; rather the Calu-3 stopped growing, rounded up and mottled, as compared to the uninfected control. The morphological change observed in Calu-3 might represent a cellular mechanism to control viral infection and therefore requires more investigation. Several other studies did not align with our findings as the strain from Munich (Zecha et al., 2020) and Japan (Matsuyama et al., 2020) showed very poor susceptibility and the Hong Kong strain showed a high susceptibility with no cytopathogenicity (Chu et al., 2020; Hui et al., 2020). The discrepant susceptibility could be dependent on the polarization of the Calu-3 cells where ACE2, the receptor for SARS-CoV-2, is expressed apically in polarized Calu-3 cells and has been shown to facilitate entry and release of the SARS-CoV (Tseng et al., 2005). Caco2 was another cell line that showed varied level of susceptibility in different studies. In Caco2 while we observed high viral susceptibility, we did not observe any appreciable cytopathogenicity as was reported for the French (Wurtz et al., 2020) and the Honk Kong strain (Chu et al., 2020). Bojkova et al. have performed a time-course proteomic study in with the Frankfurt strain of SARS-CoV-2 in infected Caco2 cells over a period of 24h (Bojkova et al., 2020). Contrary to our results (Figure 1), the authors observed cytopathogenicity in Caco2 cells 24hpi. In order to compare their 24hpi proteomic data with ours we re-analyzed their data with similar statistical considerations as ours. Compared to ours, they observed a very high number of proteins to be significantly altered (1379 vs 177), among which we observed only 30 proteins overlapping in both studies (Figure 7A). A heatmap of the 30 common proteins over time in the Bojkova et al. data is shown in Figure 7B and the heatmap of the same proteins in our study at 24hpi is shown in Figure 7C. We observed discordance in 4 proteins, where TTR and IFI35 were upregulated in their study but downregulated in ours and ITGB4 and LYPD3 were downregulated in their study but were upregulated in ours. We also specifically looked into the proteins related to IFN-signaling pathways and observed several nuclear transporters to be upregulated in the Bojkova et al. study. Among the ISGs, only ISG15 showed an upregulation in both the data (Supplemental Figure S9). Of note, unlike others, the Frankfurt strain was the only strain that was isolated and adapted in Caco2 that could have possibly led to higher susceptibility and CPE of this strain in the cell-line (Bojkova et al., 2020). In addition, different infective doses, structural changes in the virus, culture conditions and clonal differences in the cell lines could also govern strain-specific differences in cellular tropism of SARS-CoV-2.

**Figure 7.**
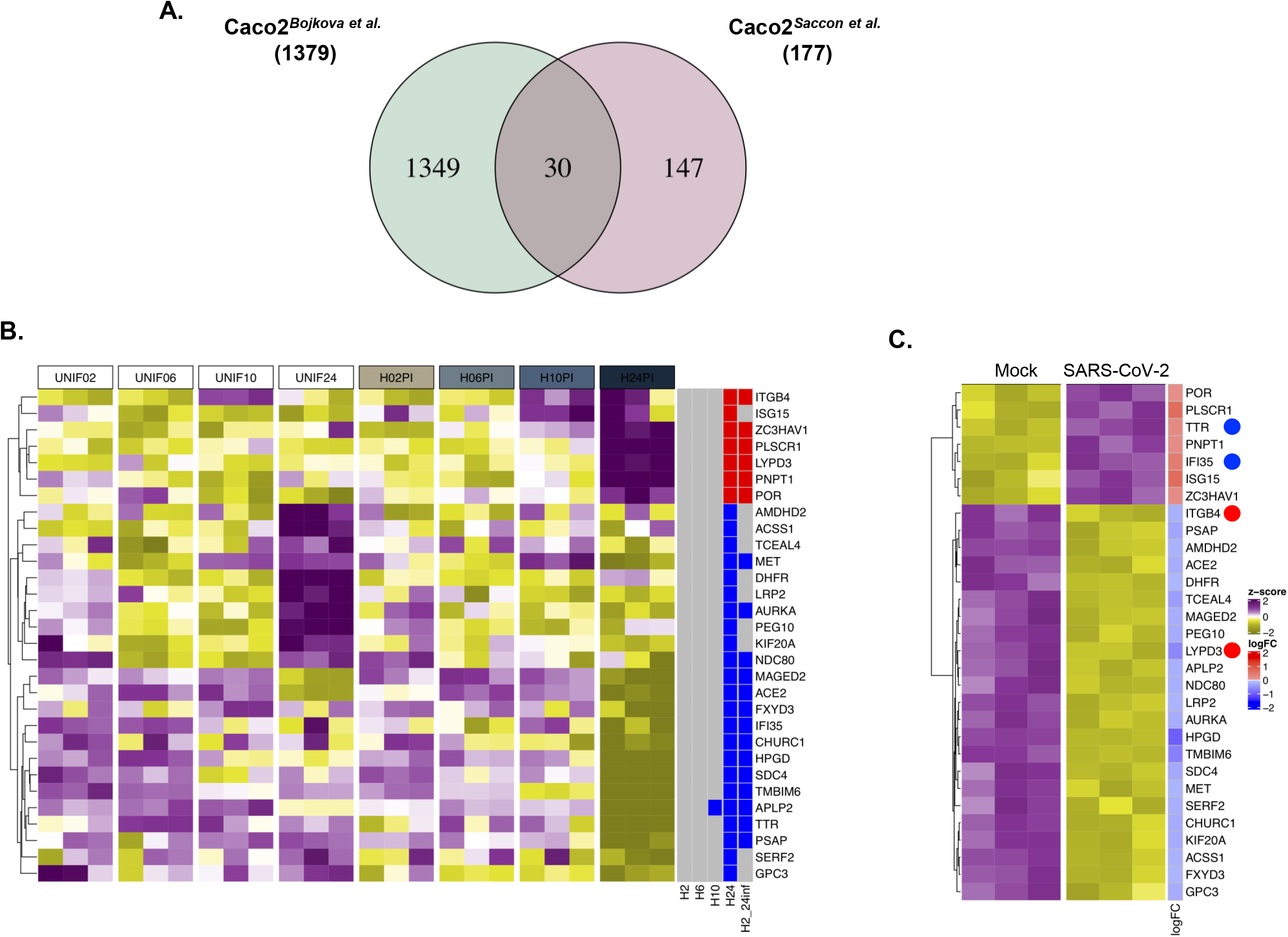
Validation of proteins dysregulated in Caco2 cell lines by SARS-CoV-2 infection from independent studies. A. Venn diagram and overlap of proteins with significant abundance between mock and 24hpi in Caco2 cell lines from this study and a study by Bojkova et al. B. Heatmap of significant overlapping proteins in mock-infected and SARS-CoV-2 infected cells at 24h in Bojkova et al. Data were quantile normalized and Z-score transformed. Lower values are represented in yellow and higher values in purple. Significant differentially expressed proteins between time points are indicated in blue if downregulated and in red if up-regulated. C. Heat map of the significant differentially abundant proteins of the Bojkova et al. in our dataset (Bojkova et al., 2020).

A major role in susceptibility to viral infection is played by the interaction between viral envelope glycoproteins and cellular receptors, which initiate the early phases of the viral life cycle. Therefore, we first investigated whether mutations in the spike glycoproteins of the different strains presented above could affect their tropism and cytopathogenicity. However, no major amino acid changes were observed in the spike protein, except for the Munich strain harboring D614G mutation that had become the dominant genotype in Europe and considered to be more infectious in humans (Korber et al., 2020). Interestingly, both Caco2 and Calu-3 were not susceptible to the Munich strain in the study performed by Zecha et al. and this correlated with the absence of ACE2 expression in their steady-state proteomics data obtained from Caco2 and Calu-3 cell lines (Zecha et al., 2020). ACE2 and TMPRSS2 proteins are considered essential for SARS-CoV-2 entry and ACE2 expression alone strongly correlated with the virus susceptibility. We observed a strong correlation between ACE2 expression and virus tropism, while increased expression of TMPRSS2 alone in Huh7 and 293FT cell lines showed susceptibility to SARS-CoV-2, perhaps due to TMPRSS2 mediated enhanced virus uptake into the cell (Heurich et al., 2014). However, we cannot exclude the possibility that other molecules or endocytosis mechanisms are involved in viral recognition and entry.

In addition, in the present study, we have investigated the replicative capacity of the virus by measuring the presence of newly produced virions in the culture supernatant. Another interesting observation was that we did not observe any virus budding in the cell surface of Huh7 and 293FT, in spite of increased viral RNA in the supernatant over time, differently from Caco2 and Calu-3 (Figure 2). Recently, it has been shown that beta-coronaviruses can hijack the lysosomal organelles resulting in non-lytic exocytosis of the virus (Ghosh et al., 2020). Overall our data on SARS-CoV-2 virus production, release, receptor expression and cytotoxicity highlight the cell-line specific SARS-CoV-2 life cycle and suggest different mechanisms of remodeling of the host cellular environment during infection, which we addressed by quantitative proteomics. While there were definite virus-specific proteome changes (Supplemental Figure S7), the pathways that were commonly regulated between Caco2 and Calu-3 belonged to type-I interferon signaling. It was also evident that the dynamics of the viral life cycle influences the proteomic landscape or vice versa. As, in Caco2 and Calu-3 an efficient production of the virus correlated with increased activation of RIG-I/MDA5 signaling and subsequently IFN-β and ISGs by 24hpi (Figure 4 and 5). However, this was not observed in 293FT and Huh7. In another study, we have observed that in Huh7 cells SARS-CoV-2 can inhibit type-I IFN signaling early and the dysregulation in this pathway is only observed 48hpi (unpublished data), which correlates with moderate production of the virus. In spite of differences in the dynamics of the cellular response, type-I IFN pre-sensitization of the Caco2, Calu-3 (Shuai et al., 2020) and Huh7 (unpublished data) has been shown to effectively inhibit SARS-CoV-2 infection. Additionally, many of the ISGs like IFIT1, ISG15 and DDX58 were upregulated in all the three cell lines (Figure 6). These findings suggest that even in presence of cell-type specific diversity in cellular responses, there are common pathways that could be efficiently targeted to inhibit SARS-CoV-2. This is particularly relevant since SARS-CoV-2 can infect different organs of the body (Mallapaty, 2020; Trypsteen et al., 2020).

One limitation of our study is that the analysis is restricted to cell lines, which may not be physiologically representative of the human tissue, contrary to organoids, but we still provide an overview of the complexity and variability of the interaction between SARS-CoV2 and the human cellular targets. Furthermore, we restricted our proteomics study to 24hpi and more detailed time kinetics experiments are required to elucidate better the dynamic changes occurring during infection.

In conclusion, we identify several cell lines of human origin that could be used as physiologically relevant cell models to study the biological properties of SARS-CoV-2. Type-I interferon is commonly regulated during infection in cell lines originating from lungs, colon and liver and warrants more mechanistic studies to identify factors that could be utilized to control the infection.

## Acknowledgments

We would like to thank Prof. Lena Palmberg, and Asst. Prof. Swapna Upadhyay, Karolinska Institute for providing the 16HBE cell line and Prof. Matti Sällberg for providing the 293FT cells. The authors would like to acknowledge the support from the Proteomics Biomedicum, Karolinska Institutet for LC-MS/MS analysis. The study is funded by Swedish Research Council Grants (2017-01330) to U.N., Karolinska Institute Stiftelser och Fonder (2020-02153 to S.G. and 2020-01554 to U.N.), Åke Wibergs Stiftelse (M20-0220) to S.G., Swedish research Council (2018-05766 and 2017-03126) and Innovative Medicines Initiative 2 Joint Undertaking (JU) under grant agreement no. 101005026 to A.M. JU receives support from the European Union’s Horizon 2020 research and innovation programme and EFPIA. TF acknowledges the grant received from the Swedish Cancer Society and the Swedish Research Council.

## Authors Contribution

**E.S.** designed the experiments, performed experimental work, analyzed the results and wrote the first draft of the manuscript. **X.C.** performed experimental work, analyzed the results and reviewed and edited the manuscript. **F.M.** performed the bioinformatics analysis and wrote the first draft of the manuscript. **Á.V.** and **J.E.R.** performed the mass-spectrometry and reviewed and edited the manuscript. **L.S.** performed the electron microscopy. **B.S.V.** and **S.K.** performed experimental work and reviewed and edited the manuscript. **T.F.** and **S.N.B** .provided intellectual content and reviewed and edited the manuscript. **A.M.** contributed with resources, intellectual content and reviewed and edited the manuscript. **U.N.** conceptualized, acquired funding, provided intellectual content and reviewed and edited the manuscript. **S.G.** conceptualized, supervised and performed the experimental work, analyzed the results, acquired funding, and wrote the manuscript together with all the co-authors. All authors have read and agreed to the published version of the manuscript.

## Data availability

All data generated or analyzed during this study are included in this published article. Information regarding any additional data are available from the corresponding author on reasonable request.

