## Supplementary Methods and Figure legends for "Tropism of SARS-CoV-2 in commonly used laboratory cell lines and their proteomic landscape during infection"

### Supplemental information

### Transparent methods

#### Cells and viruses

The hepatocyte derived cellular carcinoma cell-line Huh7 was obtained from Marburg Virology Lab (Germany) matching the STR reference profile of Huh7 (Rohde et al., 2019), Vero-E6 (ATCC® CRL-1586™), Calu-3 (ATCC#HTB-55), A549 (ATCC#CCL-185) were obtained from ATCC (USA) and Caco2 cells was obtained from CLS cell line services, GmbH, Germany (#300137). 16HBE was obtained from Lena Palmberg, Karolinska Institutet (Sweden) and 293FT (Invitrogen, #10769564) was obtained from Matti Sällberg, Karolinska Institutet (Sweden). The SARS-CoV-2 virus (SWE/01/2020) used in this study was isolated from a nasopharyngeal sample of a patient at Public Health Agency (Sweden) and the virus was confirmed as SARS-CoV-2 by genome sequencing (Genbank accession number MT093571).

#### Virus propagation and infection

SARS-CoV-2 was propagated in Vero-E6 and titrated by measuring the tissue culture infectious dose (TCID50) in Vero-E6 cells. For virus susceptibility, the different cell lines were seeded at a concentration of 10,000 cells/well in a 96-well plate, 24h prior to infection. Calu-3 cells were incubated either for 24h or 72h before infection. Infection was performed by incubating the cells with 100µL of DMEM (Sigma-Aldrich, Sweden) supplemented with 5% heat-inactivated FBS (ThermoFisher, Sweden) containing SARS-CoV-2 either at moi 1 or 0.1 for 1h at 37°C. Then, the medium was removed and replenished with fresh medium. The cells were incubated for 120hpi.

#### Virus production and cytotoxicity

The virus production in cell supernatant and the virus mediated cytotoxicity were determined at 3h, 24h, 48h, 72h, 96h and 120h post infection. The cytotoxicity was determined by measuring the cellular ATP by using Viral ToxGlo assay (Promega, Sweden) as per manufacturers guidelines. The virus RNA in the supernatant was determined by qRT-PCR targeting either the N-gene or the E-gene using Takara PrimeDirect probe, RT-qPCR mix (Takara Bio Inc, Japan) using the primers (Eurofins, USA) and probes (Sigma-Aldrich, UK) reported in supplemental table S1. RT-qPCR was performed as previously reported (Corman et al., 2020), with cycling conditions: initial denaturation 90°C for 3 min, reverse transcription 60°C for 5 min, followed by

45 cycles of 95°C for 5 sec, 58°C for 30 sec. Relative quantification of viral copies was done by comparing to the serially diluted stock virus. The cytotoxicity and virus production data for the Vero-E6, 16HBE and Huh7 at moi 1 for the time points 3hpi, 24hpi, 48hpi and 72hpi was extracted from our previously published data (Appelberg et al., 2020).

Supplemental Table S1:

| **Name** | **SARS-CoV-2 E gene** |
|---|---|
| **Fwd Sequence (5' to 3')** | ACAGGTACGTTAATAGTTAATAGCGT |
| **Rev Sequence (5' to 3')** | ATATTGCAGCAGTACGCACACA |
| **Probe Sequence (5' to 3')** | [FAM] ACACTAGCCATCCTTACTGCGCTTCG [BBQ650] |
| **Name** | **SARS-CoV-2 N gene** |
| **Fwd Sequence (5' to 3')** | CACATTGGCACCCGCAATC |
| **Rev Sequence (5' to 3')** | GAGGAACGAGAAGAGGCTTG |
| **Probe Sequence (5' to 3')** | [FAM] ACTTCCTCAAGGAACAACATTGCCA [BBQ650] |

**Quantitative RT-PCR**

Messanger RNA (mRNA) expression of a few ISG transcripts and human GAPDH were measured by qRT-PCR. The sequences of the qPCR primers are listed in supplemental table S2. Total RNA was extracted using Direct-zol™ RNA miniprep (Zymo Research, USA) and RNA concentration was assessed using a spectrophotometer (NanoDrop UV Visible Spectrophotometer, Thermofisher, USA). Reverse transcription was performed using a high capacity reverse transcription kit (Applied Biosystems, USA) for 10 min at 25°C, followed by 37°C for 120 min and 85°C for 5 min. Quantitative RT-PCR assays were setup using the Power SYBR Green PCR Master Mix (Applied Biosystems, UK) using 250nM of primer pairs with cycling conditions: initial denaturation 95°C for 10 min, followed by 40 cycles of 95°C for 15 sec, 60°C for 1 min. Melting curves were run by incubating the reaction mixtures at 95°C for 15 sec, 60°C for 20 sec, 95°C for 15 sec, ramping from 60°C to 95°C in 1°C/sec. The values were normalized to endogenous GAPDH. Fold change was calculated as: Fold Change = 2-Δ(ΔCt) where ΔCt = Ct target—Ct housekeeping and Δ(ΔCT) = ΔCt infected - ΔCt mock-infected/untreated, according to the Minimum Information for Publication of Quantitative Real-Time PCR Experiments (MIQE) guidelines.

Supplemental Table S2

| **Name** | **Fwd**<br>**Sequence (5' to 3')** | **Rev**<br>**Sequence (5' to 3')** |
|---|---|---|
| IFN-B | TCCAAATTGCTCTCCTGTTG | GCAGTATTCAAGCCTCCCAT |
| IFIT1 | TCTCAGAGGAGCCTGGCTAA | TGACATCTCAATTGCTCCAGA |
| MX1 | CCAGCTGCTGCATCCCACCC | AGGGGCGCACCTTCTCCTCA |
| MX2 | CAGAGGCAGCGGAATCGTAA | TGAAGCTCTAGCTCGGTGTTC |
| ISG15 | CGCAGATCACCCAGAAGATCG | TTCGTCGCATTTGTCCACCA |
| RIG-I | ATCCCAGTGTATGAACAGCAG | GCCTGTAACTCTATACCCATGTC |
| GAPDH | TGGGCTACACTGAGCACCAG | GGGTGTCGCTGTTGAAGTCA |

**Antibodies and chemicals**

Mouse anti-ACE2 E11, (SC390851, 1:1000), mouse anti-TMPRSS2 H4, mouse anti-ISG15 (1:1000, sc-166755) (sc-515727, 1:1000) were purchased from Santa Cruz Biotechnology Ltd (USA). Mouse-anti-β-Tubulin clone TUB2.1, (T5201, 1:1000) and mouse anti- β-actin (1:5000; A5441) were purchased from Sigma-Aldrich (USA). Rabbit anti-RIG-I clone D14G6 (1:1000; #3743), rabbit anti-MDA5 clone D74E4 (1:1000; #5321), rabbit anti-p-STAT1 (1:1000; #9167) and rabbit anti-pIRF-3 clone 4D4G (1:1000; #4947) were purchased from Cell-Signaling Technologies (USA). Rabbit SARS-CoV-2 nucleocapsid (1:1000; #BSV-COV-AB-04) was from Bioserv (UK). DL-Dithiothreitol (DTT, D0632) was purchased from Sigma-Aldrich (USA). Complete protease inhibitor cocktail and phosphatase inhibitor cocktail were purchased from Roche Diagnostic (Germany). TBS buffer was purchased from VWR (Sweden). The Tris-HCl pH7.4, NaCl, EDTA and Tween-20 stock solution were obtained from Karolinska Institutet substrate department (Sweden).

**Western Blot**

Following 24hpi and 48hpi infection with different doses of SARS-CoV-2, the cells were lysed in 2% SDS lysis buffer (50 mM Tris-Cl pH 7.4, 150 mM NaCl, 1 mM EDTA, 2% SDS, freshly supplemented with 1 mM DTT, 1x protease inhibitor cocktail and 1x phosphatase inhibitor cocktail) followed by boiling at 95°C for 10 min to inactivate the virus. The protein concentration was evaluated by DC Protein Assay from Bio-Rad (USA). Evaluation of protein expression was performed by running 20µg of total protein lysate on NuPage Bis Tris 4%-12% gels (Invitrogen, USA). Proteins were transferred using iBlot dry transfer system (Invitrogen, USA) and blocked for 1h using 5% milk or BSA in 0.1% TBS-t (0.1% Tween-20). Subsequent antibody incubations were performed at 4°C overnight or for 1h at room temperature followed by incubation for 1h at room temperature with Dako polyclonal goat anti-rabbit or anti-mouse immunoglobulins/HRP (Agilent Technologies, USA). Membranes were washed using 0.1% TBS-T and proteins were detected using ECL or ECL Select (GE Healthcare, USA) on ChemiDoc XRS+ System (Bio-Rad Laboratories, USA). The Western blot analysis was performed by using antibodies targeting ACE2, TMPRSS2, β-Tubulin, RIG-I, MDA-5, TRIM25, ISG15, p-IRF3, p-STAT1 and GAPDH.

**Scanning electron microscopy (SEM)**

SEM for SARS-CoV-2 infected cells were performed as decribed previously (Szekely et al., 2020). Briefly, Cells grown on Thermanox™ coverslips (Thermo Fischer Scientific, Sweden) were fixed by immersion in 2.5% glutaraldehyde in 0.1M phosphate buffer, pH 7.4. The coverslips were rinsed with 0.1M phosphate buffer and Milli-Q® water prior to stepwise ethanol dehydration and critical-point-drying using carbon dioxide in an EM CPD 030 (Leica Microsystems, Germany). The coverslips were mounted on alumina specimen stubs using carbon adhesive tabs and sputter coated with a thin layer of platinum using a Q150T ES (Quorum Technologies, United Kingdom). SEM images were acquired using an Ultra 55 field emission scanning electron microscope (Carl Zeiss Microscopy GmbH, Germany) at 3kV and the SE2 detector.

**Quantitative proteomics analysis**

Proteomics pipeline was performed similarly as we reported previously (Appelberg et al., 2020). Cell lines were divided in two different batches, batch 1: Calu-3 and Caco2 cells, and batch 2: 293FT and Huh7 cells. For each cell line in each batch, three biological replicates of mock-infected and SARS-CoV-2-infected cells (24h) were included. Briefly, proteins were extracted with SDS-based buffer, digestion was performed on S-Trap micro columns (Protifi, Huntington, USA) and resulting peptides were labeled with isobaric TMTpro™ reagents. Labeled peptides were fractionated by high pH (HpH) reversed-phase chromatography and concatenated to a total of 12 fractions, of which each fraction was analyzed on an Ultimate 3000 UHPLC (ThermoFisher Scientific, USA) in a 120 min linear gradient. Data were acquired on a Orbitrap™ Q-Exactive HF-X™ mass spectrometer (ThermoFisher Scientific, USA) in data dependent acquisition (DDA) mode, isolating the top 20 most intense precursors at 120,000 mass resolution in the mass range of *m/z* 375 – 1400, and applying maximum injection time (IT) of 50 ms and dynamic exclusion of 30 s; precursor isolation width of 0.7 Th with high collision energy (HCD) of 34% at resolution of 45,000 and maximum IT of 86 ms in MS2 event.

Proteins were searched against both SwissProt human and SARS-CoV2 databases using the search engine Mascot Server v2.5.1 (MatrixScience Ltd, UK) in Proteome Discoverer v2.4 (ThermoFisher Scientific, USA) software environment allowing maximum two missed cleavages. Oxidation of methionine, deamidation of asparagine and glutamine, TMTpro modification of lysine and N-termini were set as variable modifications, while carbamidomethylation of cysteine was used as fixed modification. The false discovery rate (FDR) was set to 1%.

The raw mass spectrometric data was deposited to the ProteomeXhanger Consortium (http://proteomecentral.proteomexchange.org) via the PRIDE partner repository with the dataset identifier PXD023760.

**Statistical analysis**

Statistical analyses for proteomics and transcriptomics were performed in R package LIMMA. All other statistical calculations were performed in GraphPad Prism (Version 8.0.0) using unpaired t test. Significance values are indicated in the figures and figure legends. p*<0.05, **<0.01, ***<0.001 and ****<0.0001.

**Bioinformatics analysis**

Proteomics data of SARS-CoV-2 infected (moi 1) Huh7 (Appelberg et al., 2020) and Caco2 cells (Bojkova et al., 2020) were re-analyzed for this study. Differential abundance analysis was performed using R package LIMMA between mock and 48hpi for Huh7 and respectively mock and infected cells at 0, 6, 10, 24hpi. Pairwise comparisons were extracted and Benjamini-Hochberg (BH) adjustment was applied on p values. Genes with adjusted p values <0.05 were selected. Huh7, Caco2, Calu-3 and 293FT SARS-CoV-2 infected and mock infected cells were collected at 24h and TMT-labeled proteomics was performed. Proteomics raw data was first filtered for empty rows and quantile normalized with R package NormalizerDE. Histogram was used to display the distribution of data and assess that the distribution follows a normal law. Principal component analysis was performed using ggplot2 (supplemental table S3). Viral protein abundances were retrieved and baseline subtraction (Infected-Mock) was performed for each time point and represented using barplots made with ggplot2. Differential abundance analysis was performed using LIMMA as described previously. Three manually curated libraries based on interferon-regulated genes were created based on reactome terms "Antiviral mechanism by IFN−stimulated genes", "Interferon gamma signaling" and "Interferon alpha/beta signaling" (https://reactome.org/). Each library had respectively 89, 172 and 138 genes. The total number of interferon-regulated genes excluding overlap between libraries is 205 (supplemental table S4).

Supplemental Table S3

| **Cell line** | **Detected** | **DPA** |
|---|---|---|
| 293FT | 8977 | 0 |
| Huh7 (new) | 7623 | 4 |
| Caco2 | 8784 | 177 |
| Calu-3 | 8768 | 6462 |
| Munch | 6258 | 1448(24H) |
| Huh7 (old) | 8991 | 3830 |

Supplemental Table S4

| Cell line | Detected (IFN) | DPA (IFN) |
|---|---|---|
| 293FT | 109 | 0 |
| Huh7 (new) | 109 | 0 |
| Caco2 | 131 | 27 |
| Calu-3 | 129 | 105 |
| Munch | 93 | 17(24H) |
| Huh7 (old) | 97 | 46 |

Protein profiles were represented as a heatmap using R ComplexHeatmap function. Venn diagrams were made using interactivenn (http://www.interactivenn.net/). Dotplot was made using ggplot2. Interferon-regulated significant proteins (LIMMA, FDR <0.05) were represented as a network with Cytoscape ver 3.6.1. For each node, fold changes were added to the network template file. Protein-protein interactions were retrieved from STRING Db (v5.0) (https://string-db.org/). Interactions were filtered on confidence score with minimum interaction of 0,700. Only interactions from databases and experiences were conserved.

R package ReactomePA was used for reactome pathway-based analysis. Pathways analysis results were represented using barplot enrichment maps available from the package. To compare biological pathways among several cell lines, R package clusterProfiler was used.

**Supplementary figures**

**Figure S1:** A. Venn diagram of proteins with higher abundance in Caco2 and Calu-3 cell lines after SARS-CoV-2 infection compared to mock. B. Venn diagram of proteins with lower abundance in Caco2 and Calu-3 cell lines after SARS-CoV-2 infection compared to mock.

**Figure S2**: Heatmaps representing the number of significant proteins (LIMMA, FDR < 0.05) between mock and 24hpi in A. 293FT and B. Huh7 cell lines.

**Figure S3**. Cytoscape networks of differentially abundant IFN-stimulated proteins in A. Calu-3 and B. Caco2 cell lines. Proteins are represented as circles. Gradient color was applied on proteins depending on fold change (low = blue to high = red). Size of protein is proportional to the fold change.

**Figure S4**. Expression levels of A. ACE2 receptor and B. TMPRSS2 receptor in Caco2 and Calu-3 cells upon infection at 24hpi as quantified by TMT-based proteomics.

**Figure S5**. A. ISG15 protein levels in SARS-CoV-2 infected or mock-infected cells. The representative western blot with indicated antibodies are shown. B. The intensity of specific bands was quantified by ImageJ and fold change was calculated relative to the mock-infected cells, normalized to β-actin.

**Figure S6.** A. Venn diagram of proteins with higher abundance in Caco2, Calu-3 and Huh7 cell lines after 24h of SARS-CoV-2 infection compared to mock. B. Venn diagram of proteins with lower abundance in Caco2, Calu-3 and Huh7 cell lines after 24h of SARS-CoV-2 infection compared to mock.

**Figure S7**. A. ReactomePA barplot enrichment map of significant proteins identified in Calu-3 cell line excluding Caco2 and Huh7. B. ReactomePA barplot enrichment map of significant proteins identified in Huh7 cell line excluding Caco2 and Calu-3.

**Figure S8**. A. ReactomePA barplot enrichment map of significant overlapping proteins in Huh7 and Caco2 cell lines. B. ReactomePA barplot enrichment map of significant overlapping proteins in Huh7 and Calu-3 cell lines.

**Figure S9.** Re-analysis of data from Bojkova et al. paper showing heatmap representing the number of significant proteins (LIMMA, FDR $< 0.05$) belonging to interferon-signaling pathways between mock and 24hpi in Caco2 cell line.
