## Supplementary figures and images for "Tropism of SARS-CoV-2 in commonly used laboratory cell lines and their proteomic landscape during infection"

### Supplementary Figures

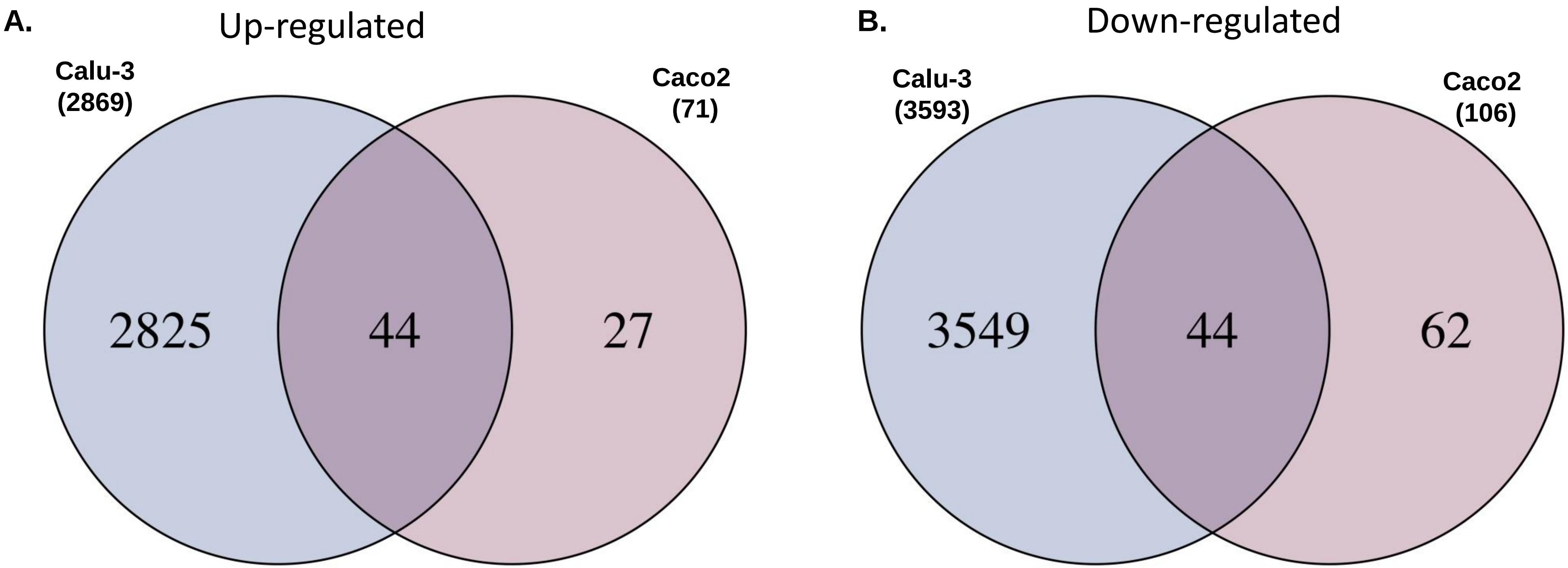

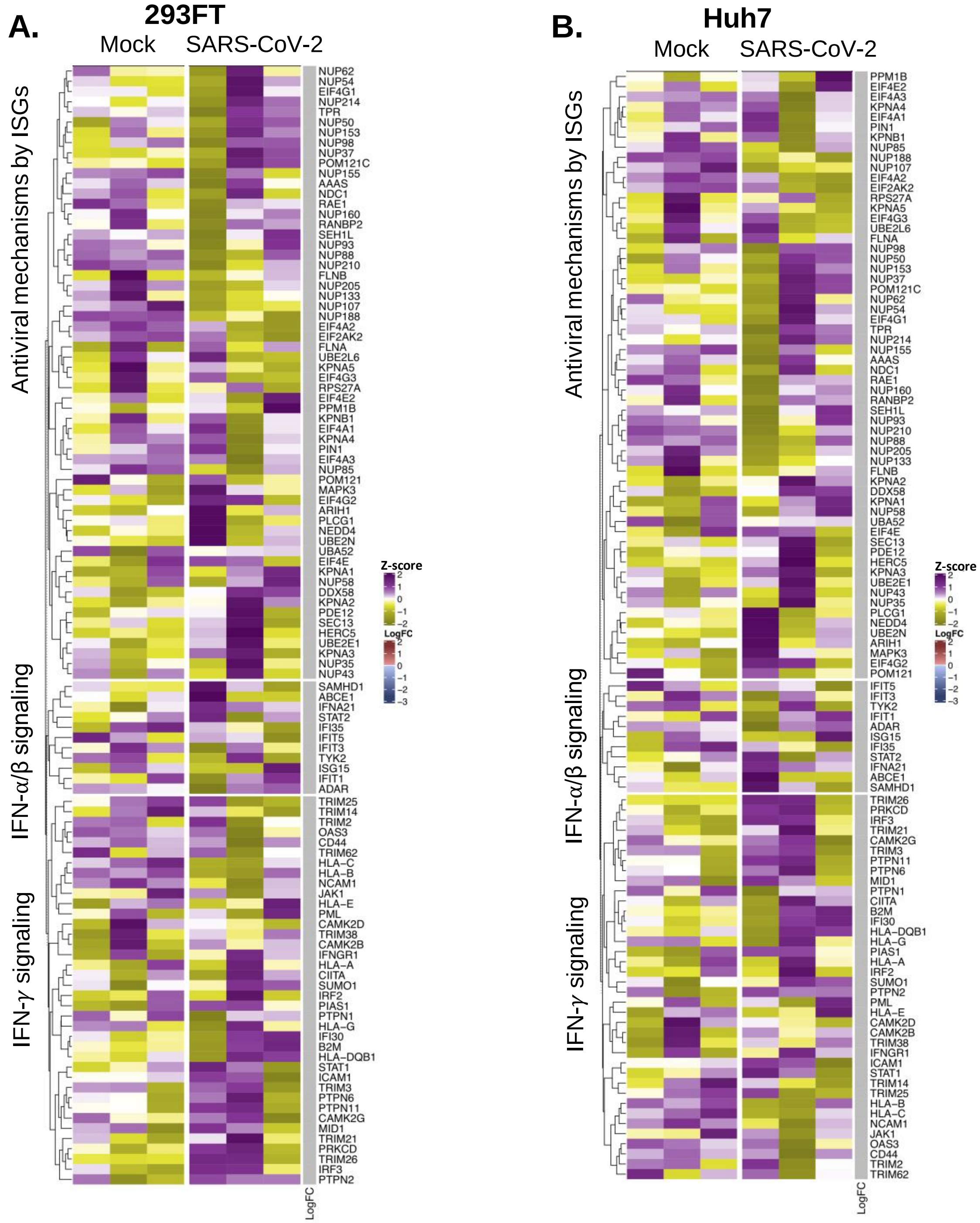

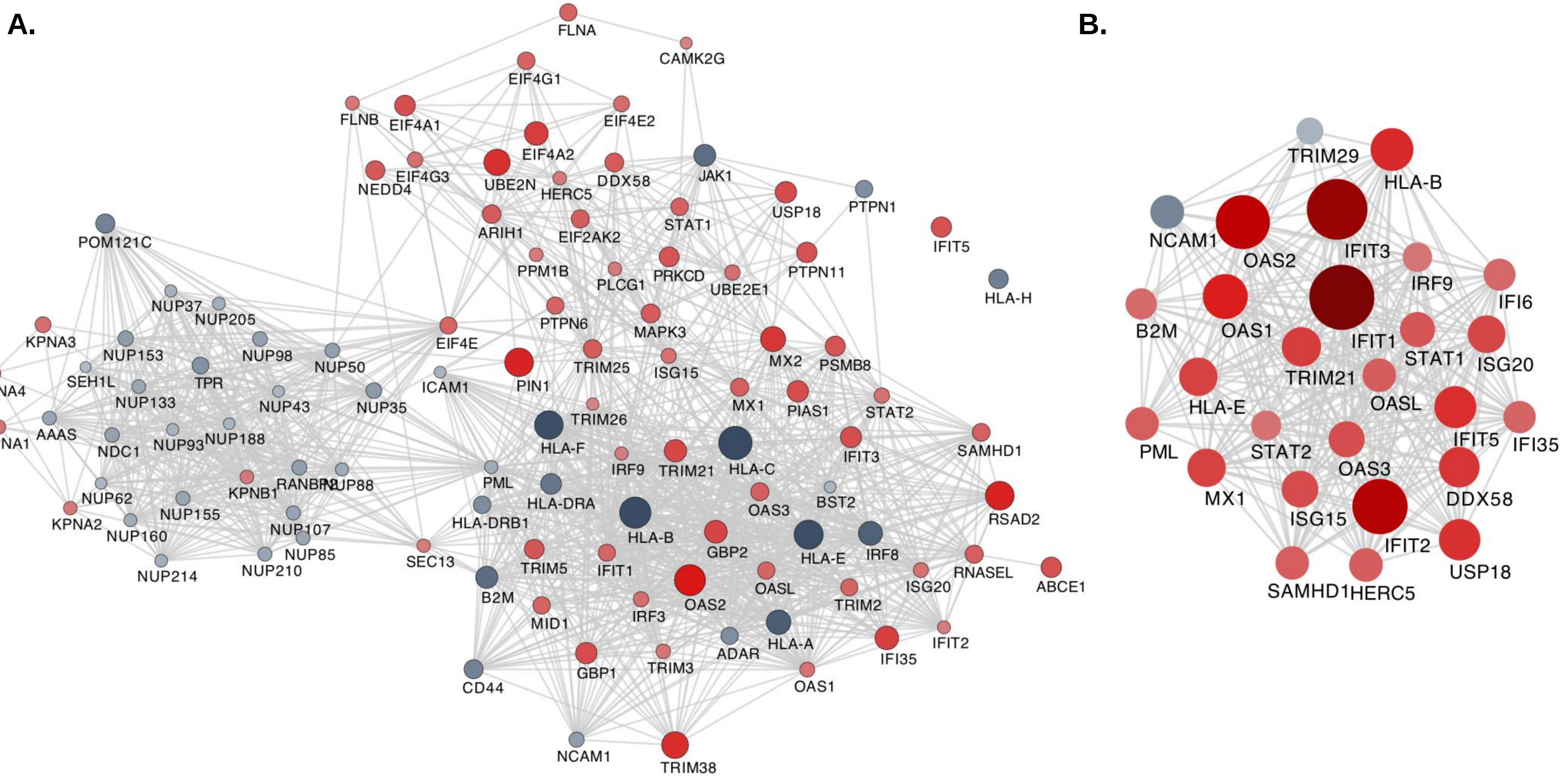

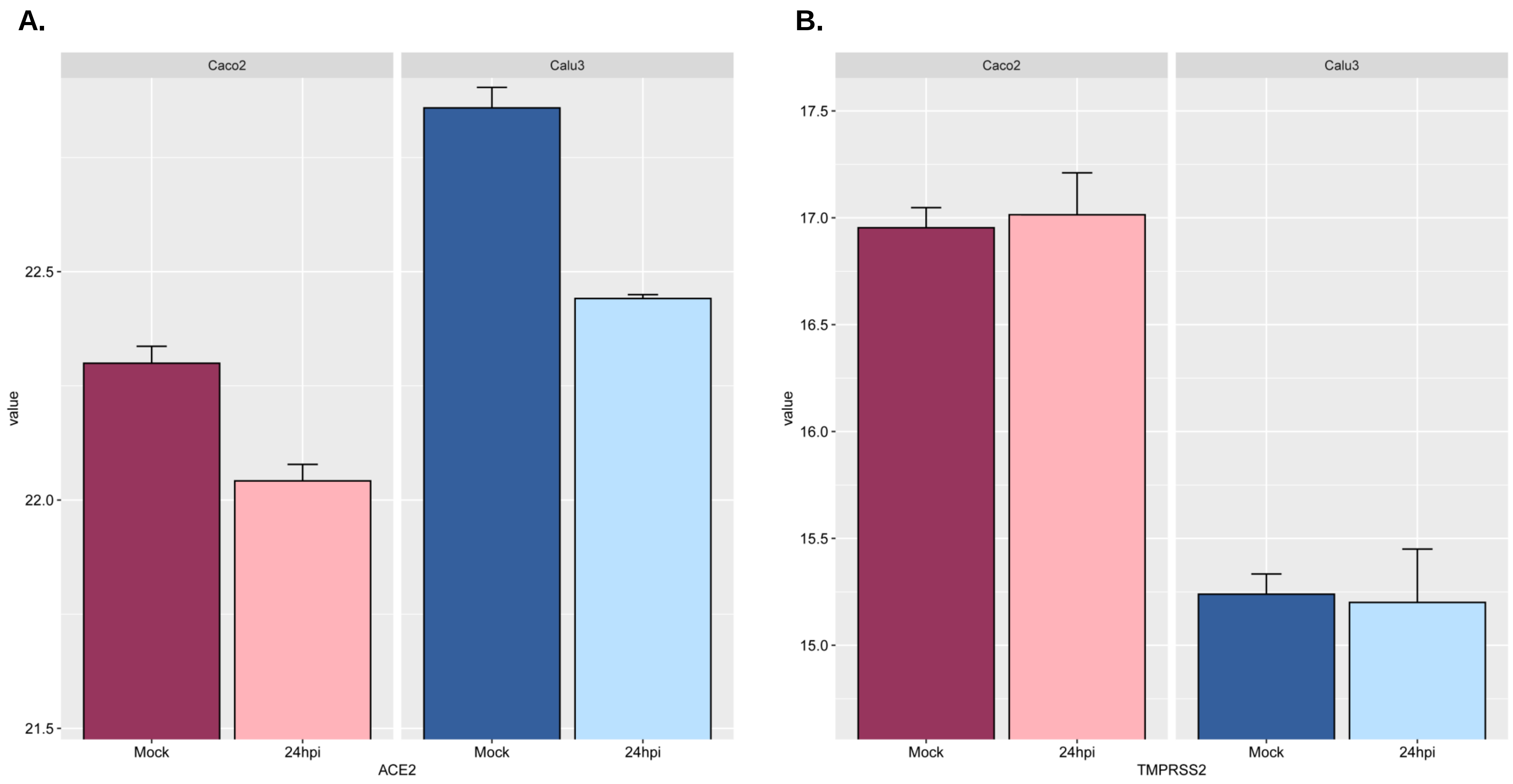

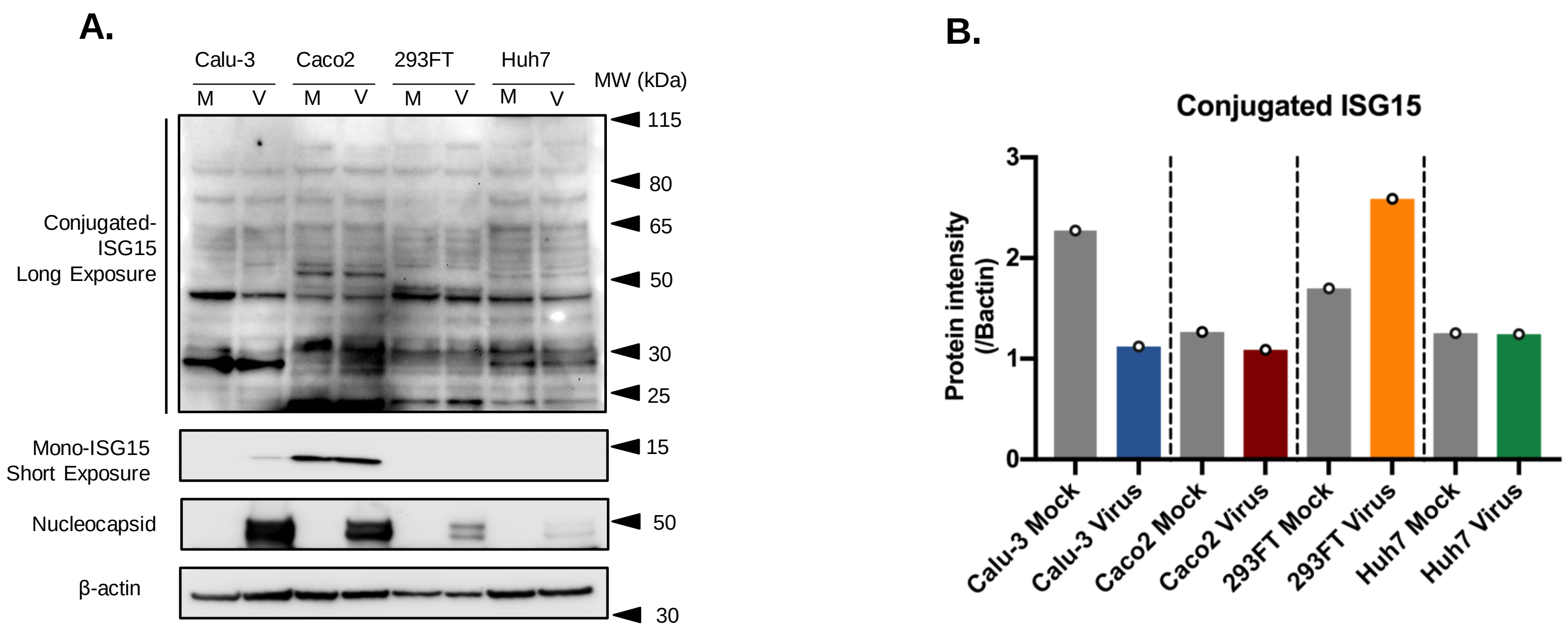

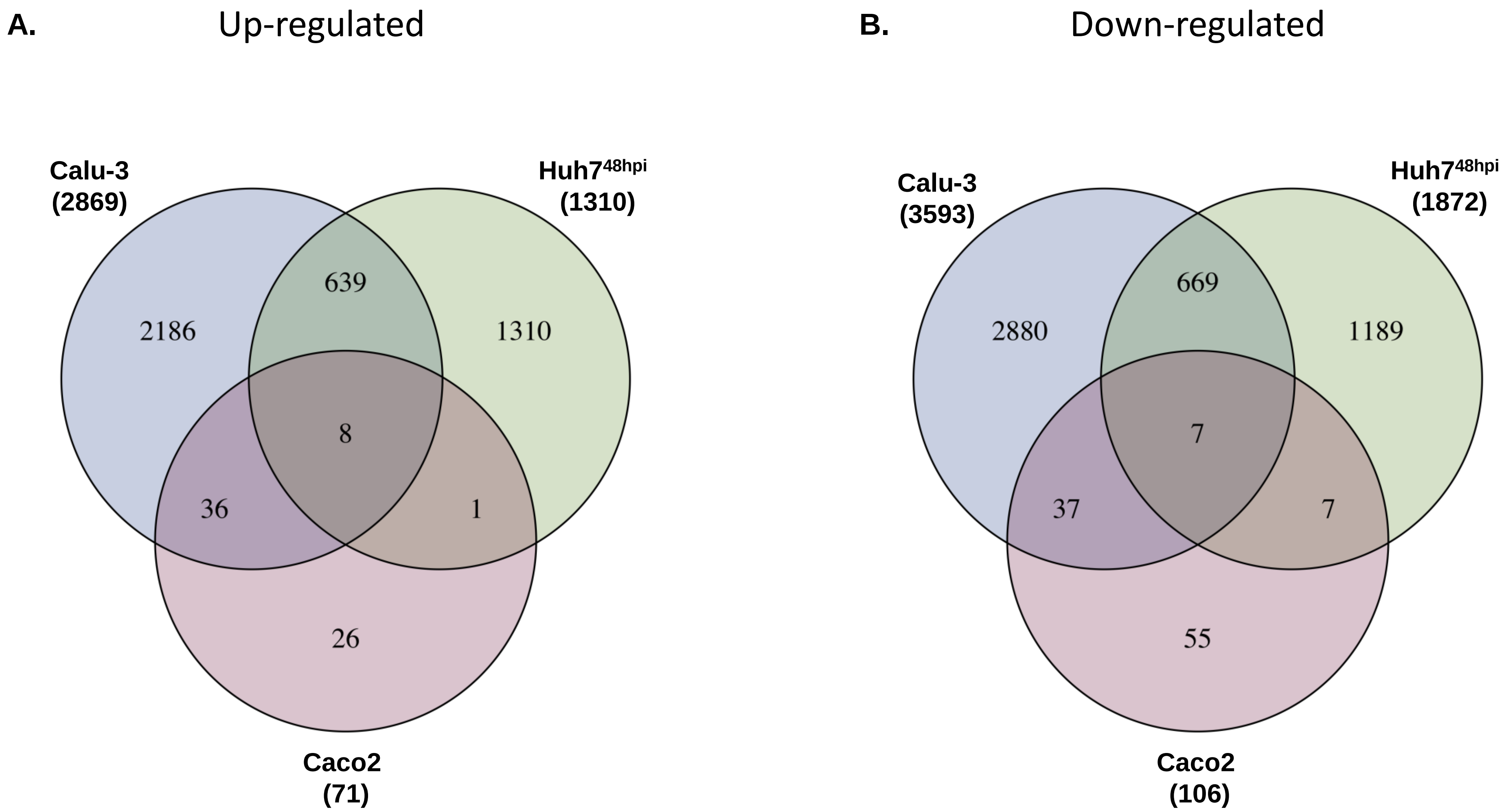

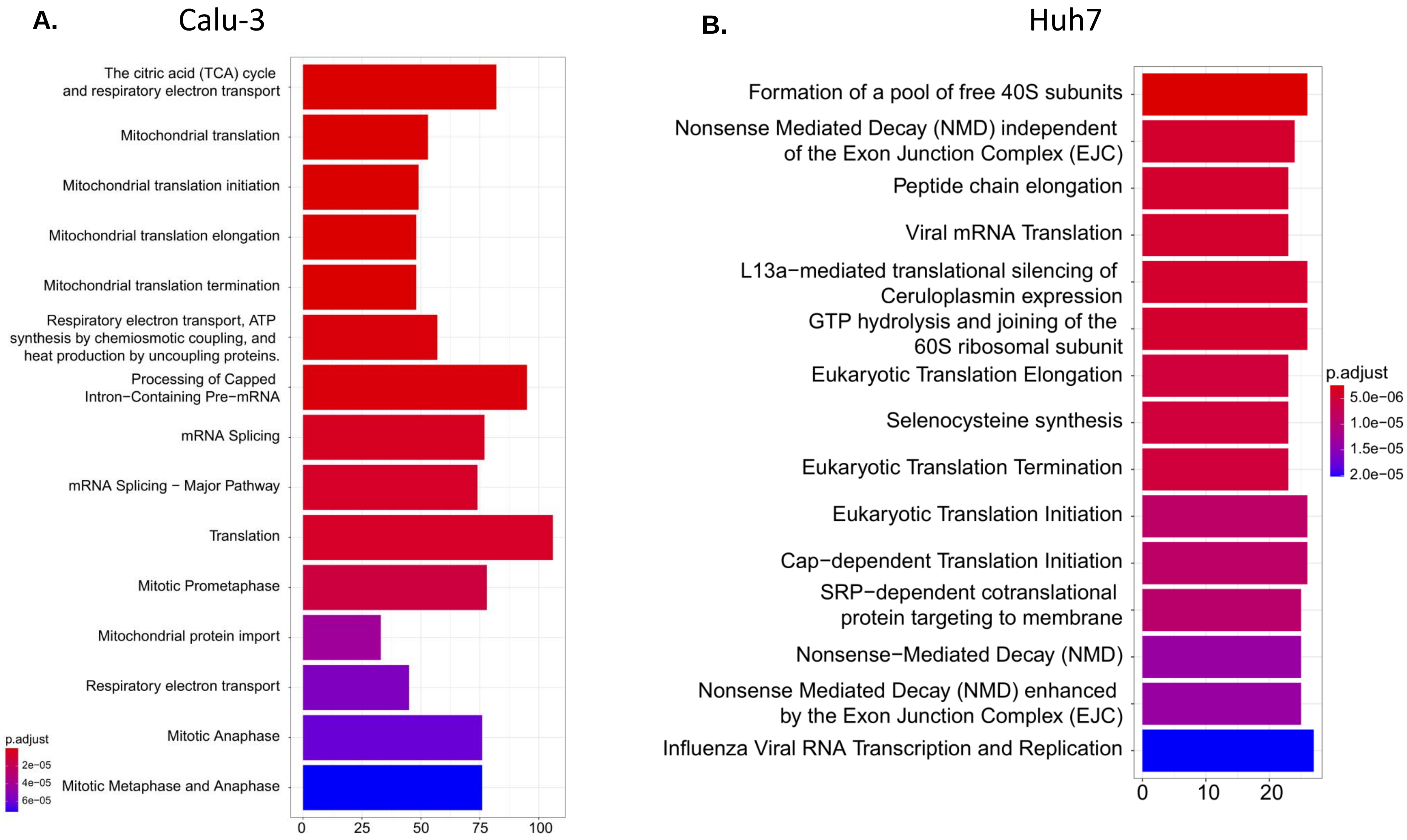

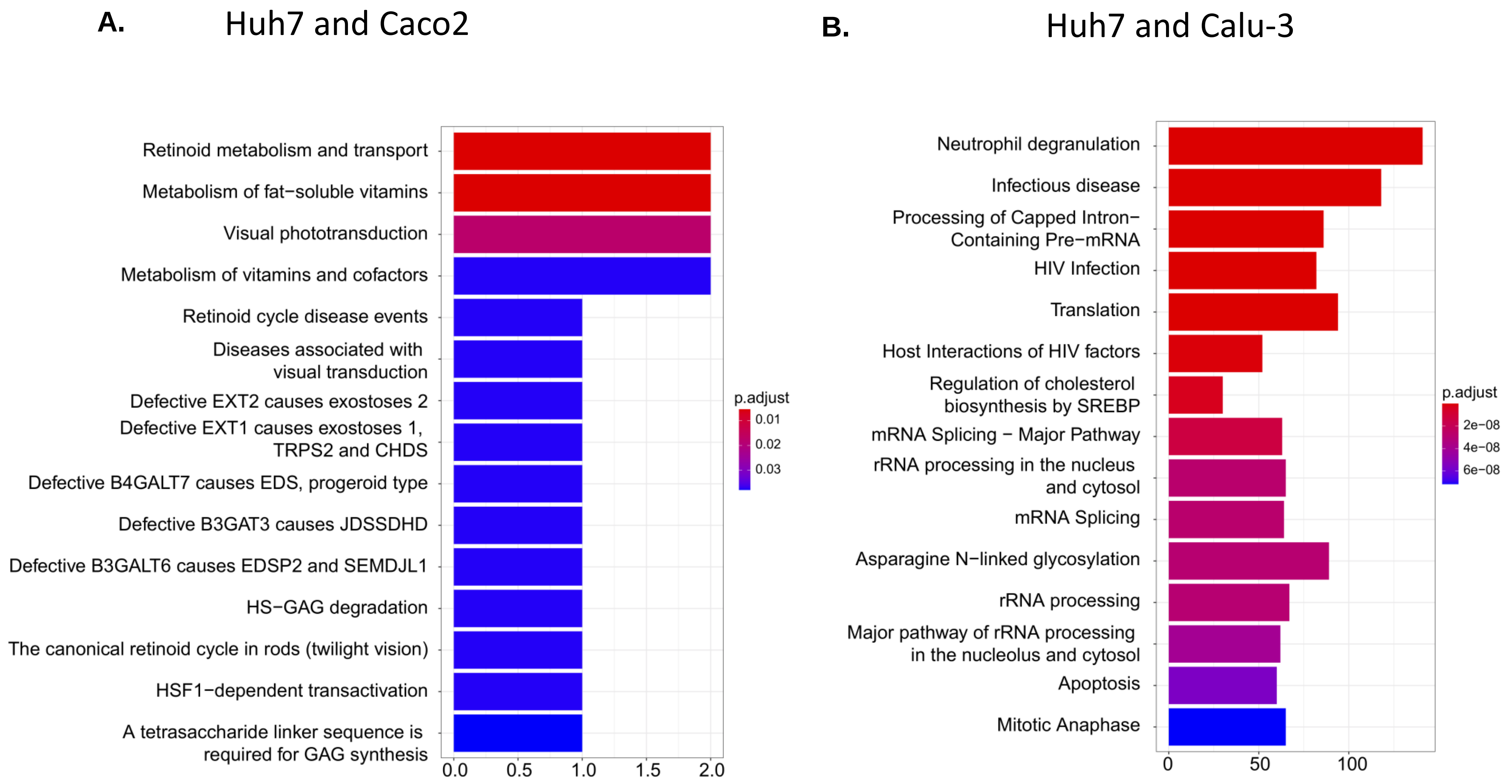

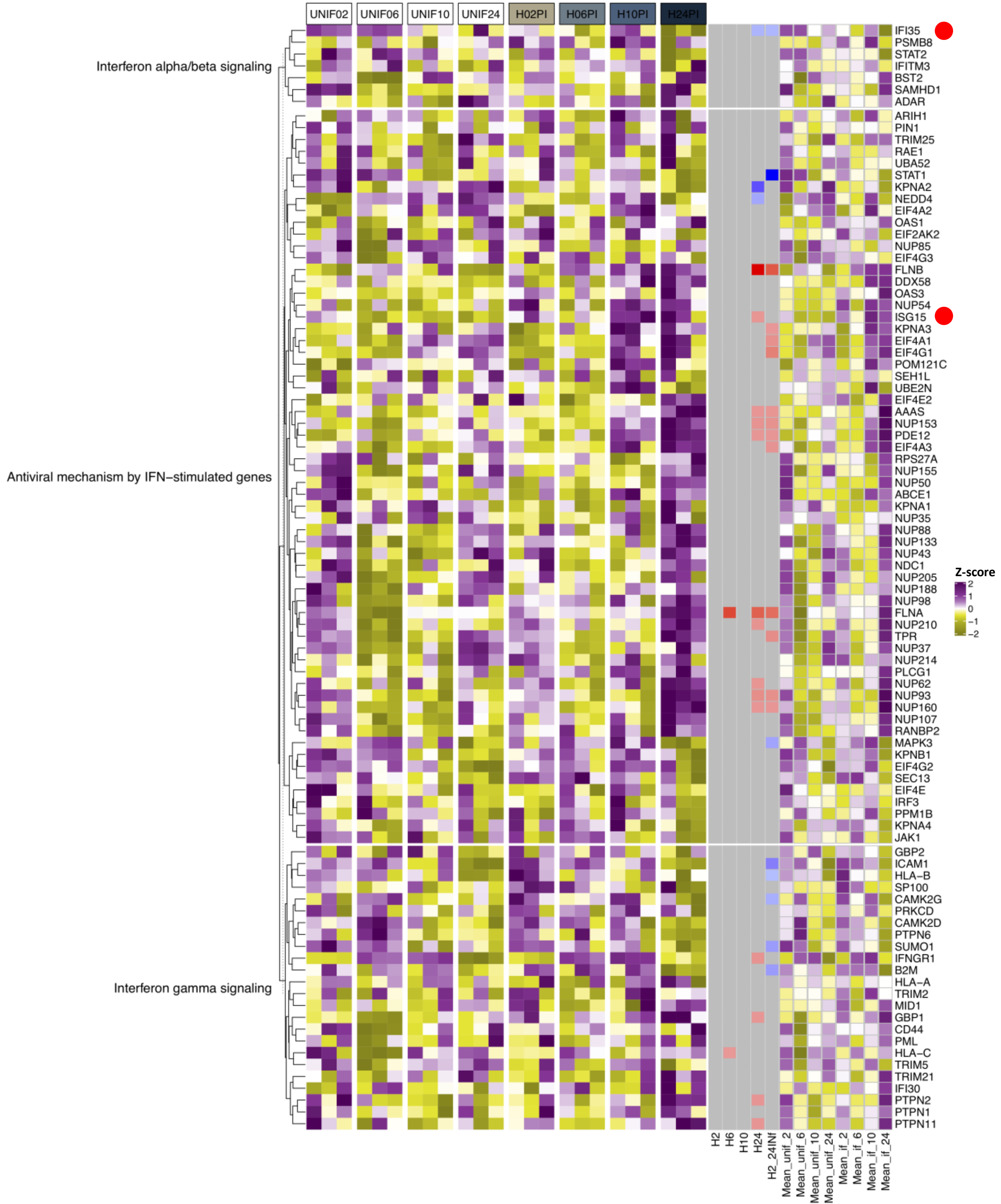
